# Mechanistic Insights into Sex Differences in Atrial Electrophysiology and Arrhythmia Vulnerability through Sex-specific Computational Models

**DOI:** 10.1101/2025.08.18.670886

**Authors:** Nathaniel T. Herrera, Haibo Ni, Charlotte E. R. Smith, Yixuan Wu, Dobromir Dobrev, Stefano Morotti, Eleonora Grandi

## Abstract

Atrial fibrillation (AF), the most common cardiac arrhythmia, is characterized by notable sex differences in clinical presentation, treatment response, and outcomes. Although prevalence is similar between sexes, women often experience more severe symptoms, higher rates of adverse drug effects, and reduced treatment efficacy. To investigate the underlying sex-specific AF mechanisms, we developed and validated male and female human atrial cardiomyocyte models that integrate known sex-based differences in electrophysiology and calcium (Ca^2+^) handling under normal sinus rhythm (nSR) and chronic AF (cAF) conditions. While the model parameterizations were based on limited human data, and the assumptions may not capture the full spectrum of clinical variability, the models reproduced key reported sex-dependent differences in human atrial cardiomyocyte action potential (AP) and Ca^2+^ transient (CaT) dynamics. Simulations revealed that both sexes exhibited shortened effective refractory periods and wavelengths in cAF vs. nSR. However, females were more prone to delayed afterdepolarizations (DADs), while males were more susceptible to AP duration (APD) and CaT amplitude (CaT_Amp_) alternans. Population-based modeling identified distinct parameter associations with arrhythmia mechanisms, whereby DAD vulnerability was associated with enhanced ryanodine receptor sensitivity to Ca^2+^ (in females), and alternans in males correlated with reduced L-type Ca^2+^ current maximal conductance. Pharmacological simulations revealed sex-specific responses to antiarrhythmic therapies. In males, multiple drug combinations proved effective in restoring APD at 90% repolarization (APD_90_), CaT_Amp_, and reducing alternans susceptibility, whereas females responded to only one combination improving APD_90_ and CaT_Amp_ but with minimal impact on DAD risk. These findings underscore the need for sex-specific therapeutic strategies and support the use of computational modeling in guiding precision medicine approaches against AF.

**Key points:**

- Atrial fibrillation (AF) is a common heart rhythm disorder that presents differently in males and females, but how the underlying mechanisms differ in males and females is not fully understood.
- We developed and validated computer models of male and female human atrial cardiomyocytes that incorporate known sex differences in ion channels and calcium handling under normal sinus rhythm and AF conditions.
- Under normal rhythm, males and females showed distinct electrical activity, which became less pronounced in AF. In AF, both sexes showed reduced effective refractory period and wavelength and depressed calcium transients. Males were more susceptible to electrical alternans, while females showed a greater tendency for calcium-driven delayed afterdepolarizations.
- Simulated drug treatments showed greater benefit in male models, particularly with combinations targeting multiple potassium channels, while female models showed limited response. These results highlight the need for sex-specific approaches to treating AF and may help guide future drug development.

## Introduction

Atrial fibrillation (AF), is the most common cardiac arrhythmia worldwide and is projected to triple in prevalence over the next 50 years, representing a growing global epidemic (Kornej *et al*., 2020). AF is characterized by a rapid and irregular heartbeat, which can lead to palpitations, dyspnea, and chest pain (Rienstra *et al*., 2012), and is associated with an increased risk of death due to stroke, myocardial infarction, and heart failure (Streur, 2019). While men are more likely to develop AF at a younger age, the overall prevalence is similar in women due to their longer life expectancy (Ko *et al*., 2016; Odening *et al*., 2019). Yet, despite similar prevalence, women often present with more comorbidities (Khaing *et al*., 2025), experience more severe and persistent symptoms, and have a higher risk of adverse thrombo-embolic outcomes, which results in reduced quality of life (Ball *et al*., 2013; Ko *et al*., 2016; Odening *et al*., 2019).

Variation in treatment efficacy also exists between sexes in AF. Women are referred five-times less frequently for catheter ablation compared to men, and when treated, they experience lower ablation success rates (Patel *et al*., 2010). They also have greater sensitivity to antiarrhythmic drugs like amiodarone (Essebag *et al*., 2007), and a higher incidence of adverse effects following pharmacological treatment often leading to treatment discontinuation (Muzzey *et al*., 2020).

These disparities in clinical presentation, treatment response, and outcomes suggest underlying biological differences in AF pathophysiology between sexes. Recent research has highlighted important sex differences in cardiac electrophysiology (Gaborit *et al*., 2010), yet much of the existing work has focused on the ventricles. In the atria, emerging evidence suggests that sex-based differences exist not only in baseline electrophysiology (EP) and calcium (Ca^2+^) handling properties during normal sinus rhythm (nSR), but also in the nature and extent of remodeling that occurs with the onset and progression of AF to a chronic form (cAF, Smith, Ni and Grandi, 2025). These differences span ion channel and Ca^2+^ cycling protein densities, gene expression profiles, and regulatory signaling pathways (Ambrosi *et al*., 2013; Herraiz-Martínez *et al*., 2022; Smith, Ni and Grandi, 2025) and likely contribute to sex-specific variations in action potential (AP, Ravens, 2018; Pecha *et al*., 2023) and Ca^2+^ transient (CaT) dynamics, as well as differential susceptibility to arrhythmia (Zhang *et al*., 2024). However, at present the mechanisms remain incompletely understood.

Understanding sex-differences in atrial electrophysiological remodeling and arrhythmia susceptibility is essential for developing more effective, personalized strategies for AF prevention and treatment. We recently developed a sex-specific human atrial model focused on subcellular Ca^2+^ handling mechanisms, which recapitulated the experimentally observed increase in arrhythmogenic Ca^2+^ release events in females with AF (Herraiz-Martínez *et al*., 2022) and identified potential Ca^2+^ targeted therapeutic interventions (Zhang *et al*., 2024). However, existing computational models have not comprehensively incorporated sex-based differences in atrial EP. Zhang *et al*. (2024) utilized AP clamp to characterize arrhythmogenic Ca^2+^ release, thus preventing a comprehensive assessment of the bidirectional coupling between transmembrane voltage and Ca^2+^. In the present study, we expand on our prior work by integrating both EP and Ca^2+^ handling differences into detailed sex-specific human atrial cardiomyocyte models under nSR and cAF conditions. Model simulations predicted that although both sexes exhibit shorter refractory periods and wavelengths in cAF compared to nSR, females are more likely to develop delayed afterdepolarizations (DADs), while males show a greater propensity for alternans. These findings suggest that the mechanisms underlying AF initiation and maintenance (and/or recurrence) may differ in males and females. Using a population-based modeling approach we identify key model parameters that contribute to sex-specific differences in atrial electrophysiological properties and arrhythmia risk, and tested how potential therapeutic targets may differentially benefit males and females.

## Methods

### Parameterization of the male and female models

In this study, we expanded our recently developed model of human atrial cardiomyocyte EP and Ca^2+^ handling (Ni *et al*., 2023) to incorporate sex-differences experimentally observed in nSR and cAF. The baseline model represents a male atrial cardiomyocyte in nSR. To better align with experimental data from human atrial appendages (Pecha *et al*., 2023), we incorporated a 20% reduction in the maximal conductance of the inward rectifier K^+^ current (G_K1_) into the baseline male model. Using this adjusted baseline, we developed an analogous female atrial cardiomyocyte model under nSR by reparametrizing ion channel, ion transport, and Ca^2+^ handling properties based on experimentally observed sex differences, as summarized in **Table 1**. We then extended both the male and female nSR models to cAF conditions. We fitted the parameters of the cAF atrial male cardiomyocyte model using a parameter optimization routine to recapitulate experimentally observed AP (Pecha *et al*., 2023) and CaT properties (Heijman *et al*., 2020). This routine employed the Nelder-Mead algorithm (Nelder and Mead, 1965) to minimize the error function between simulated and experimental results, identifying the optimal set of parameter scaling factors for the cAF cellular model. Targets of model optimizations are reported in **Table 2** and initial estimates were based on our previous cAF parameterization (Grandi *et al*., 2011; Herrera *et al*., 2023). These parameter scaling factors were then applied to the female cAF atrial cardiomyocyte model. In female cAF, we did not include aspects of male cAF remodeling, such as lower phospholamban (PLB) expression and reduced I_CaL_ density and slowed inactivation kinetics, and added elevated RyR phosphorylation (Herraiz-Martínez *et al*., 2022). It is important to note that multivariate regression analysis accounting for confounding factors revealed no significant interactions between AF, sex or AF plus sex and I_CaL_ density (Herraiz-Martínez *et al*., 2022), but pairwise comparisons showed a significant reduction in I_CaL_ density in male but not in female patients with AF (Herraiz-Martínez *et al*., 2022). Furthermore, for other parameters sample size was insufficient for multivariate analyses. Therefore, while we simulated differential male to female cAF remodeling as a hypothesis-generating assumption, it requires additional experimental confirmation in larger patient cohorts.

**Table 1.**
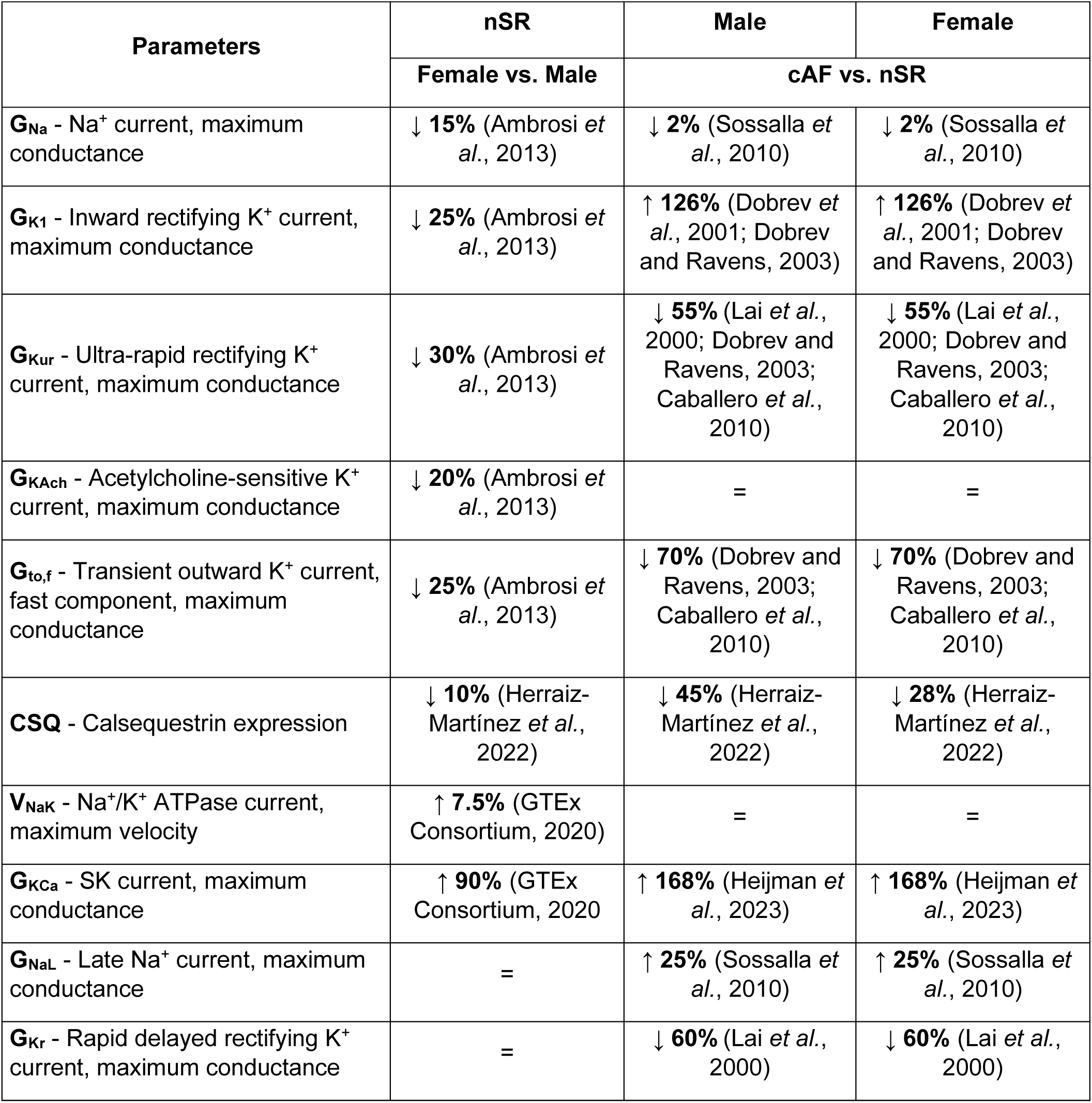
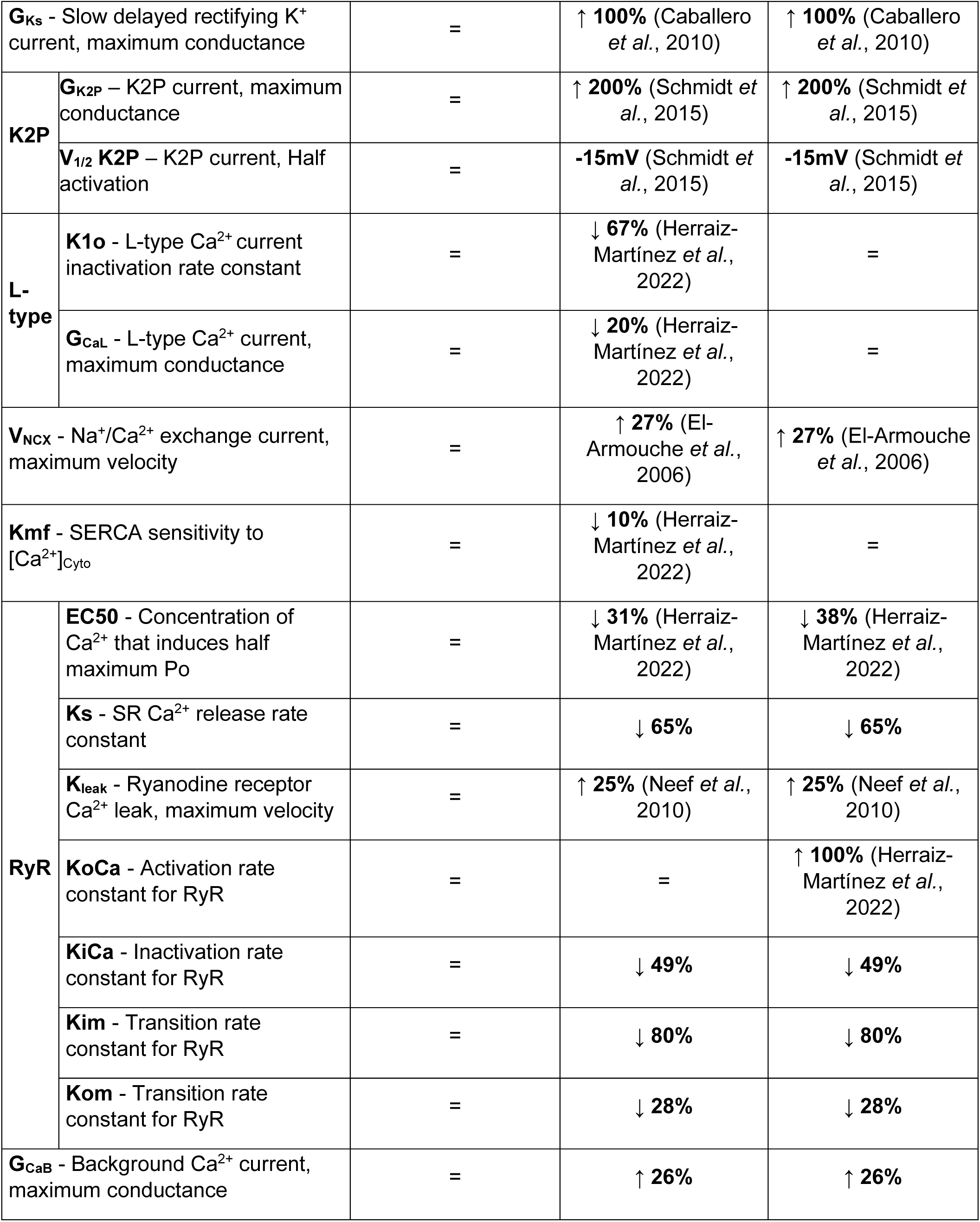
Sex- and cAF-dependent changes in electrophysiology and Ca^2+^ handling processes. Model parameters varied in the baseline male model (Ni *et al*., 2023) were adjusted to construct the female nSR model (Ambrosi *et al*., 2013; GTEx Consortium, 2020). Male cAF model parameters were optimized to match biomarker values listed in Table 2 (Lai *et al*., 2000; Brundel *et al*., 2001; Dobrev *et al*., 2001; Dobrev and Ravens, 2003; El-Armouche *et al*., 2006; Caballero *et al*., 2010; Neef *et al*., 2010; Sossalla *et al*., 2010; Schmidt *et al*., 2015; Herraiz-Martínez *et al*., 2022; Heijman *et al*., 2023), and the resulting scaling factors were subsequently applied to the female nSR model to generate the female cAF model.

| Parameters | nSR | Male | Female |
| --- | --- | --- | --- |
|  | Female vs. Male | cAF vs. nSR |  |
| $\mathbf{G_{Na}}$ - $\text{Na}^+$ current, maximum conductance | ↓ **15%** (Ambrosi *et al.*, 2013) | ↓ **2%** (Sossalla *et al.*, 2010) | ↓ **2%** (Sossalla *et al.*, 2010) |
| $\mathbf{G_{K1}}$ - Inward rectifying $\text{K}^+$ current, maximum conductance | ↓ **25%** (Ambrosi *et al.*, 2013) | ↑ **126%** (Dobrev *et al.*, 2001; Dobrev and Ravens, 2003) | ↑ **126%** (Dobrev *et al.*, 2001; Dobrev and Ravens, 2003) |
| $\mathbf{G_{Kur}}$ - Ultra-rapid rectifying $\text{K}^+$ current, maximum conductance | ↓ **30%** (Ambrosi *et al.*, 2013) | ↓ **55%** (Lai *et al.*, 2000; Dobrev and Ravens, 2003; Caballero *et al.*, 2010) | ↓ **55%** (Lai *et al.*, 2000; Dobrev and Ravens, 2003; Caballero *et al.*, 2010) |
| $\mathbf{G_{KAch}}$ - Acetylcholine-sensitive $\text{K}^+$ current, maximum conductance | ↓ **20%** (Ambrosi *et al.*, 2013) | = | = |
| $\mathbf{G_{to,f}}$ - Transient outward $\text{K}^+$ current, fast component, maximum conductance | ↓ **25%** (Ambrosi *et al.*, 2013) | ↓ **70%** (Dobrev and Ravens, 2003; Caballero *et al.*, 2010) | ↓ **70%** (Dobrev and Ravens, 2003; Caballero *et al.*, 2010) |
| $\mathbf{CSQ}$ - Calsequestrin expression | ↓ **10%** (Herraiz-Martínez *et al.*, 2022) | ↓ **45%** (Herraiz-Martínez *et al.*, 2022) | ↓ **28%** (Herraiz-Martínez *et al.*, 2022) |
| $\mathbf{V_{NaK}}$ - $\text{Na}^+/\text{K}^+$ ATPase current, maximum velocity | ↑ **7.5%** (GTEx Consortium, 2020) | = | = |
| $\mathbf{G_{KCa}}$ - SK current, maximum conductance | ↑ **90%** (GTEx Consortium, 2020) | ↑ **168%** (Heijman *et al.*, 2023) | ↑ **168%** (Heijman *et al.*, 2023) |
| $\mathbf{G_{NaL}}$ - Late $\text{Na}^+$ current, maximum conductance | = | ↑ **25%** (Sossalla *et al.*, 2010) | ↑ **25%** (Sossalla *et al.*, 2010) |
| $\mathbf{G_{Kr}}$ - Rapid delayed rectifying $\text{K}^+$ current, maximum conductance | = | ↓ **60%** (Lai *et al.*, 2000) | ↓ **60%** (Lai *et al.*, 2000) |

|  |  |  |  |  |
| --- | --- | --- | --- | --- |
| **G<sub>Ks</sub>** - Slow delayed rectifying K<sup>+</sup> current, maximum conductance |  | = | ↑ **100%** (Caballero *et al.*, 2010) | ↑ **100%** (Caballero *et al.*, 2010) |
| **K2P** | **G<sub>K2P</sub>** – K2P current, maximum conductance | = | ↑ **200%** (Schmidt *et al.*, 2015) | ↑ **200%** (Schmidt *et al.*, 2015) |
|  | **V<sub>1/2</sub> K2P** – K2P current, Half activation | = | **-15mV** (Schmidt *et al.*, 2015) | **-15mV** (Schmidt *et al.*, 2015) |
| **L-type** | **K<sub>1o</sub>** - L-type Ca<sup>2+</sup> current inactivation rate constant | = | ↓ **67%** (Herraiz-Martínez *et al.*, 2022) | = |
|  | **G<sub>CaL</sub>** - L-type Ca<sup>2+</sup> current, maximum conductance | = | ↓ **20%** (Herraiz-Martínez *et al.*, 2022) | = |
| **V<sub>NCX</sub>** - Na<sup>+</sup>/Ca<sup>2+</sup> exchange current, maximum velocity |  | = | ↑ **27%** (El-Armouche *et al.*, 2006) | ↑ **27%** (El-Armouche *et al.*, 2006) |
| **K<sub>mf</sub>** - SERCA sensitivity to [Ca<sup>2+</sup>]<sub>Cyto</sub> |  | = | ↓ **10%** (Herraiz-Martínez *et al.*, 2022) | = |
| **RyR** | **EC50** - Concentration of Ca<sup>2+</sup> that induces half maximum Po | = | ↓ **31%** (Herraiz-Martínez *et al.*, 2022) | ↓ **38%** (Herraiz-Martínez *et al.*, 2022) |
|  | **K<sub>s</sub>** - SR Ca<sup>2+</sup> release rate constant | = | ↓ **65%** | ↓ **65%** |
|  | **K<sub>leak</sub>** - Ryanodine receptor Ca<sup>2+</sup> leak, maximum velocity | = | ↑ **25%** (Neef *et al.*, 2010) | ↑ **25%** (Neef *et al.*, 2010) |
|  | **KoCa** - Activation rate constant for RyR | = | = | ↑ **100%** (Herraiz-Martínez *et al.*, 2022) |
|  | **KiCa** - Inactivation rate constant for RyR | = | ↓ **49%** | ↓ **49%** |
|  | **Kim** - Transition rate constant for RyR | = | ↓ **80%** | ↓ **80%** |
|  | **Kom** - Transition rate constant for RyR | = | ↓ **28%** | ↓ **28%** |
| **G<sub>CaB</sub>** - Background Ca<sup>2+</sup> current, maximum conductance |  | = | ↑ **26%** | ↑ **26%** |

**Table 2.** Parameters and targets used in the cost function for optimization of the male cAF model. The Nelder-Mead algorithm (Nelder & Mead, 1965) was employed to optimize the male cAF model parameters to align with experimental electrophysiological data at 1 Hz pacing (Pecha *et al*., 2023) and arrhythmia susceptibility measures (Narayan *et al*., 2011), as previously described (Heijman *et al*., 2023). Each feature included in the cost function is listed along with its target value and assigned weight, which reflects its relative importance.

| Feature included in cost function | Value | Weight |
| --- | --- | --- |
| Action potential duration 90% | 220ms | 1 |
| Action potential duration 50% | 100ms | 1 |
| Resting membrane potential | -82.5mV | 1 |
| Action potential amplitude | 125mV | 1 |
| Maximum upstroke velocity | 245V/s | 1 |
| Plateau voltage | -4.5mV | 0.2 |
| [Ca<sup>2+</sup>]<sub>i</sub> Max | 350nM | 2 |
| [Ca<sup>2+</sup>]<sub>i</sub> Min | 200nM | 2 |
| [Ca<sup>2+</sup>]<sub>SR,max</sub> | 0.50mM | 1 |
| DAD BCL threshold | 450-500ms | 1 |
| Alternans BCL threshold | 400-500ms | 1 |

### Validation of the male and female models

The sex-specific human atrial nSR and cAF cardiomyocyte models were validated by comparing model-derived biomarkers to experimental measurements of AP and CaT properties.

Specifically, we assessed whether each model condition could reproduce key electrophysiological and Ca^2+^ handling biomarkers, including action potential durations at 90% and 50% repolarization (APD_90_ and APD_50_), resting membrane potential (RMP), AP amplitude (APA), maximum upstroke velocity (V_max_), plateau voltage (V_PLT_), and CaT amplitude (CaT_Amp_), under 1 Hz steady-state pacing (Ravens, 2018; Heijman *et al*., 2020; Pecha *et al*., 2023). As this data was used for fitting the male cAF model, we validated the resulting model against an independent dataset of AP features across a range of frequencies. Male and female human atrial cell models were simulated under both nSR and cAF conditions, and AP and CaT biomarkers were evaluated at steady state across pacing frequencies ranging from 1 to 4 Hz (in 0.5 Hz increments).

We further constructed homogenous sex-specific nSR and cAF one-dimensional (1D) strands of 210 atrial cells to determine the effective refractory period (ERP), conduction velocity (CV), and wavelength (WL). Steady state initial conditions were established in the single cell and applied to the cable. To account for cAF-associated fibrosis, we incorporated a 40% reduction in tissue conductivity in both the male and female strands. This was achieved by adjusting the diffusion coefficient, which controls electrotonic coupling between cells, from 0.1441 mm²·ms⁻¹ in nSR to 0.0865 mm²·ms⁻¹ in cAF to yield physiologically realistic CVs, as done previously (Ni *et al*., 2020). At 1 Hz pacing, simulated CV was 0.67 m·s⁻¹ (male) and 0.63 m·s⁻¹ (female) in nSR and decreased to 0.46 m·s⁻¹ and 0.43 m·s⁻¹ in cAF, respectively, consistent with experimental observations not stratified by sex (Hansson *et al*., 1998; Krul *et al*., 2015; Silva Cunha *et al*., 2024). To calculate the ERP, the first 10 cells of the cable were stimulated for 6 beats, after which an S2 beat was applied 560 ms after the last S1, and a binary search was employed by reducing the S2 in 1 ms increments to determine the shortest S2 leading to AP propagation along the cable. The WL was subsequently calculated as the product between ERP and CV.

### Arrhythmia pacing protocols

We investigated the role of sex differences in atrial cardiomyocyte arrhythmogenesis by examining the determinants of APD and CaT_Amp_ alternans and DADs. Alternans occurrence was assessed in nSR and cAF male and female models at steady state by progressively decreasing the basic cycle length (BCL, at 1 ms decrements) and using a binary search to determine the BCL at which the beat-to-beat difference in APD (estimated at E_m_ -55 mV) is ≥ 5 ms, as previously done (Herrera *et al*., 2023). CaT_Amp_ alternans threshold was determined with a similar protocol and using a beat-to-beat difference of 50 nM. DAD occurrence was evaluated by pacing the models to steady-state in the presence of 1 µM isoproterenol, followed by a 30 s pause. The BCL pacing threshold for the development of DADs (depolarizations ≥ 10 mV during the no-stimulation period), was determined by progressively decreasing the BCL in 1 ms decrements in BCL, as done previously (Herrera *et al*., 2023).

### Populations of sex-specific human atrial cardiomyocyte models

We built populations of models to quantify the impact of parameter perturbations on AP and CaT features and pacing thresholds for APD alternans and DADs (Morotti and Grandi, 2024). We generated populations of 1,000 male and 1,000 female atrial cardiomyocytes in nSR and cAF by randomly perturbing key model parameters shown in **Table 4**. Model parameters were varied independently based on a log-normal distribution with a scaling factor of σ = 0.1 using a previously established approach (Sobie, 2009). To correlate changes in model parameters with variations in model outputs we performed linear regression analysis (Sobie, 2009). For each parameter, we calculated the regression coefficient along with its 95% confidence interval (CI) and corresponding p-value. A parameter was considered statistically significant if its 95% CI excluded zero and its p-value was <0.05. We displayed the regression coefficients of the top 20 most influential parameters for each biomarker

**Table 3.** Validation of male and female nSR and female cAF human atrial models: Model outputs for action potential duration at 90% repolarization (APD_90_), APD_50_, resting membrane potential (RMP), AP amplitude (APA), maximum upstroke velocity (V_max_), plateau voltage (V_PLT_), and Ca^2+^ transient amplitude (CaT_Amp_) compared to experimental results from (Ravens, 2018; Heijman *et al*., 2020; Pecha *et al*., 2023). Symbols indicating significant differences between groups (p < 0.05), were as reported in Pecha *et al*., 2023 and Ravens, 2018.

|  | **nSR** |  |  |  |  |  |
| --- | --- | --- | --- | --- | --- | --- |
|  | **Male** | **Female** | **M vs. F** | **Male** | **Female** | **M vs. F** |
| **Biomarker** | **Model** |  |  | **Experiment** |  |  |
| APD<sub>90</sub> (ms) | 299.69 | 291.96 | = | 319.91±2.60 | 314.80±4.19 | **N.S.** (Ravens, 2018; Pecha *et al.*, 2023) |
| APD<sub>50</sub> (ms) | 108.96 | 119.63 | < | 136.98±2.39 | 146.87±3.85 | < (Ravens, 2018; Pecha *et al.*, 2023) |
| RMP (mV) | -78.45 | -77.21 | < | -74.58±0.24 | -73.16±0.39 | < (Ravens, 2018; Pecha *et al.*, 2023) |
| APA (mV) | 114.73 | 113.83 | > | 95.50±0.47 | 92.8±0.75 | > (Ravens, 2018; Pecha *et al.*, 2023) |
| V<sub>max</sub> (V/s) | 237.02 | 204.80 | > | 234.04±4.50 | 212.07±7.27 | > (Ravens, 2018; Pecha *et al.*, 2023) |
| V<sub>PLT</sub> (mV) | -11.09 | -7.86 | < | -16.78±0.57 | -15.11±0.92 | **N.S.** (Pecha *et al.*, 2023)<br>< (Ravens, 2018) |
| CaT<sub>Amp</sub> (nM) | 259.96 | 275.21 | < | 217.11±135.64 | 261.63±175.19 | **N.S.** (Heijman *et al.*, 2020) |
|  | **cAF** |  |  |  |  |  |
|  | **Male** | **Female** | **M vs. F** | **Male** | **Female** | **M vs. F** |
| **Biomarker** | **Model** |  |  | **Experiment** |  |  |
| APD<sub>90</sub> (ms) | 230.55 | 219.13 | = | 216.01±3.34 | 222.19±4.30 | **N.S.** (Ravens, 2018; Pecha *et al.*, 2023) |
| APD<sub>50</sub> (ms) | 101.47 | 102.57 | = | 99.19±3.09 | 101.79±3.98 | **N.S.** (Ravens, 2018; Pecha *et al.*, 2023) |
| RMP (mV) | -82.51 | -82.32 | = | -77.57±0.30 | -77.27±0.39 | **N.S.** (Ravens, 2018; Pecha *et al.*, 2023) |
| APA (mV) | 125.57 | 123.42 | = | 102.20±0.59 | 100.98±0.77 | **N.S.** (Ravens, 2018; Pecha *et al.*, 2023) |
| V<sub>max</sub> (V/s) | 239.70 | 212.64 | > | 248.23±5.65 | 233.1±7.28 | **N.S.** (Pecha *et al.*, 2023)<br>> (Ravens, 2018) |
| V<sub>PLT</sub> (mV) | -5.97 | -5.83 | = | -5.56±0.72 | -6.63±0.93 | **N.S.** (Ravens, 2018; Pecha *et al.*, 2023) |
| CaT<sub>Amp</sub> (nM) | 170.53 | 203.70 | < | / | / | / |

**Table 4.** Definition of model parameters that are randomly altered to introduce variability into the population of models.

| **Model Parameters used in Sensitivity Analysis** |  |  |
| --- | --- | --- |
| **G<sub>Na</sub>** |  | Na<sup>+</sup> current, maximum conductance |
| **G<sub>K1</sub>** |  | Inward rectifying K<sup>+</sup> current, maximum conductance |
| **G<sub>Kur</sub>** |  | Ultra-rapid rectifying K<sup>+</sup> current, maximum conductance |
| **G<sub>KAch</sub>** |  | Acetylcholine-sensitive K<sup>+</sup> current, maximum conductance |
| **G<sub>to,f</sub>** |  | Transient outward K<sup>+</sup> current, fast component, maximum conductance |
| **CSQ** |  | Calsequestrin expression |
| **V<sub>NaK</sub>** |  | Na<sup>+</sup>/K<sup>+</sup> ATPase current, maximum velocity |
| **G<sub>KCa</sub>** |  | SK current, maximum conductance |
| **G<sub>NaL</sub>** |  | Late Na<sup>+</sup> current, maximum conductance |
| **G<sub>Kr</sub>** |  | Rapid delayed rectifying K<sup>+</sup> current, maximum conductance |
| **G<sub>Ks</sub>** |  | Slow delayed rectifying K<sup>+</sup> current, maximum conductance |
| **G<sub>K2P</sub>** |  | K2P current, maximum conductance |
| **L-Type** | **K1o** | L-type Ca<sup>2+</sup> current inactivation rate constant |
|  | **CaMKII-p** | CaMKII-phosphorylation of L-type Ca<sup>2+</sup> current |
|  | **G<sub>CaL</sub>** | L-type Ca<sup>2+</sup> current, maximum conductance |
| **V<sub>NCX</sub>** |  | Na<sup>+</sup>/Ca<sup>2+</sup> exchange current, maximum velocity |
| **Kmf** |  | SERCA sensitivity to [Ca<sup>2+</sup>]<sub>Cyto</sub> |
| **RyR** | **EC50** | Concentration of Ca<sup>2+</sup> that induces half maximum P<sub>o</sub> |
|  | **V<sub>RyR</sub>** | Ryanodine receptor flux, maximum velocity |
|  | **K<sub>leak</sub>** | Ryanodine receptor Ca<sup>2+</sup> leak, maximum velocity |
|  | **KoCa** | Activation rate constant for RyR |
|  | **KiCa** | Inactivation rate constant for RyR |
|  | **Kim** | Transition rate constant for RyR |
|  | **Kom** | Transition rate constant for RyR |
| **G<sub>CaB</sub>** |  | Background Ca<sup>2+</sup> current, maximum conductance |
| **G<sub>ClB</sub>** |  | Background Cl<sup>-</sup> current, maximum conductance |
| **G<sub>ClCa</sub>** |  | Ca<sup>2+</sup>-activated Cl<sup>-</sup> current, maximum conductance |
| **G<sub>NaB</sub>** |  | Background Na<sup>+</sup> current, maximum conductance |
| **G<sub>Kp</sub>** |  | Plateau K<sup>+</sup> current, maximum conductance |
| **V<sub>SERCA</sub>** |  | Sarcoplasmic reticulum Ca<sup>2+</sup> ATPase flux, maximum velocity |
| **V<sub>PMCA</sub>** |  | Plasma membrane Ca<sup>2+</sup> ATPase current, maximum velocity |
| **Ca<sup>2+</sup> Buffer Cleft** |  | Ca<sup>2+</sup> buffering strength in the cleft space |
| **Ca<sup>2+</sup> Buffer Cytosol** |  | Ca<sup>2+</sup> buffering strength in the cytosolic space |

### Drug simulations

Simulations were performed using the cAF male and female atrial cell models to assess the effects of pharmacological interventions on atrial AP, Ca^2+^ handling, and arrhythmia vulnerability. Male and female cAF models were subjected to 25% inhibition of the small-conductance Ca^2+^-activated K^+^ (SK) channel maximum conductance (G_SK_), 25% inhibition of the two-pore domain K^+^ (K2P) channel maximum conductance (G_K2P_), and combined G_SK_ and G_K2P_ inhibition. Additionally, the effects of flecainide on I_Kr_, I_Na_, and I_CaL_ were incorporated as previously described (Iseppe *et al*., 2021), across concentrations corresponding to 1x, 2x, and 3x their effective free therapeutic plasma concentrations (EFTPC). We modeled the effects on the ryanodine receptor (RyR) as previously described (Yang *et al*., 2016), by modifying the RyR state model to include a drug-bound open state (DO), with transitions governed by 𝐾_on_ = 𝐷 ∗ [𝐷𝑟𝑢𝑔] and 𝐾_off_ = 𝐷 ∗ 𝐼𝐶_SO,Drug_ relative to the open state (O), diffusion rate of flecainide of 5500M^-1^ms^-1^ and IC_50_ of 20 µM. The effects of G_SK_, G_K2P_, and G_SK_+G_K2P_ inhibition were simulated both alone and in combination with varying concentrations of flecainide.

### Numerical methods and code

Single cell simulations were performed with an Intel Core i9-10900X CPU at 3.70 GHz 10 Cores using MATLAB R2022a (MathWorks, Natick, MA), and simulations of 1D cables were performed with a computing cluster with Intel Xeon CPU E5-2690 v4 at 2.60 GHz 28 CPUs (56 threads) + 132 GB. Data analysis was performed with MATLAB. The source code of our updated nSR and cAF sex-specific human atrial cell models can be accessed at https://github.com/drgrandilab.

## Results

### Validation of sex-specific human atrial cardiomyocyte models

We developed and validated sex-specific human atrial cardiomyocyte models that integrate EP and Ca^2+^ handling under both nSR and cAF conditions (**Fig. 1*A***). These models incorporate experimentally observed sex differences in the expression of ionic and Ca^2+^ handling proteins. In nSR, females exhibit lower levels of Nav1.5, Kv4.3, KChIP2, Kir2.1, Kv1.5, and Kir3.1, and calsequestrin-2 (CSQ) compared to males, along with higher transcript levels in human atrial appendage tissue for ATP1A1, the predominant Na^+^/K^+^-ATPase isoform, and KCNN2, the predominant SK channel isoform (**Table 1**; Ambrosi *et al*., 2013; GTEx Consortium, 2020; Herraiz-Martínez *et al*., 2022). For cAF, we incorporated additional remodeling features, including lower phospholamban (PLB) expression and reduced I_CaL_ density in males, as well as elevated RyR phosphorylation in females, implemented as an increase in the RyR activation rate constant KoCa (Herraiz-Martínez *et al*., 2022). The sex-specific models were validated against multiple independent experimental datasets. In nSR, the models reproduce known sex differences (Ravens, 2018; Pecha *et al*., 2023), with females displaying longer APD_50_ and higher V_PLT_, along with a more depolarized RMP, and reduced APA and V_max_, while APD_90_ remains similar between sexes (**Table 3**). In nSR the female model exhibits a higher V_PLT_, longer APD_50_, and a lower V_max_ compared to the male model across multiple pacing frequencies. Notably, the differences in APD_50_ and V_max_ were attenuated at higher pacing rates (**Fig. 1*B-I***). In cAF, these EP differences between sexes are largely abolished, in agreement with experimental data at 1 Hz pacing (**Table 3**, Pecha *et al*., 2023; Ravens, 2018). Additionally, the simulated AP and CaT features align closely with multiple experimental datasets across a range of pacing frequencies, further validating the models. In cAF, both male and female models have a decreased APD_90_, APD_50_, CaT_Amp_, an increased V_max_, V_PLT_, and a more hyperpolarized RMP compared to their respective nSR conditions (**Fig. 1*B-I***). Of note, at higher pacing frequencies several EP and CaT features exhibit a bifurcation that is consistent with the emergence of AP and Ca^2+^-alternans.

**Figure 1.**
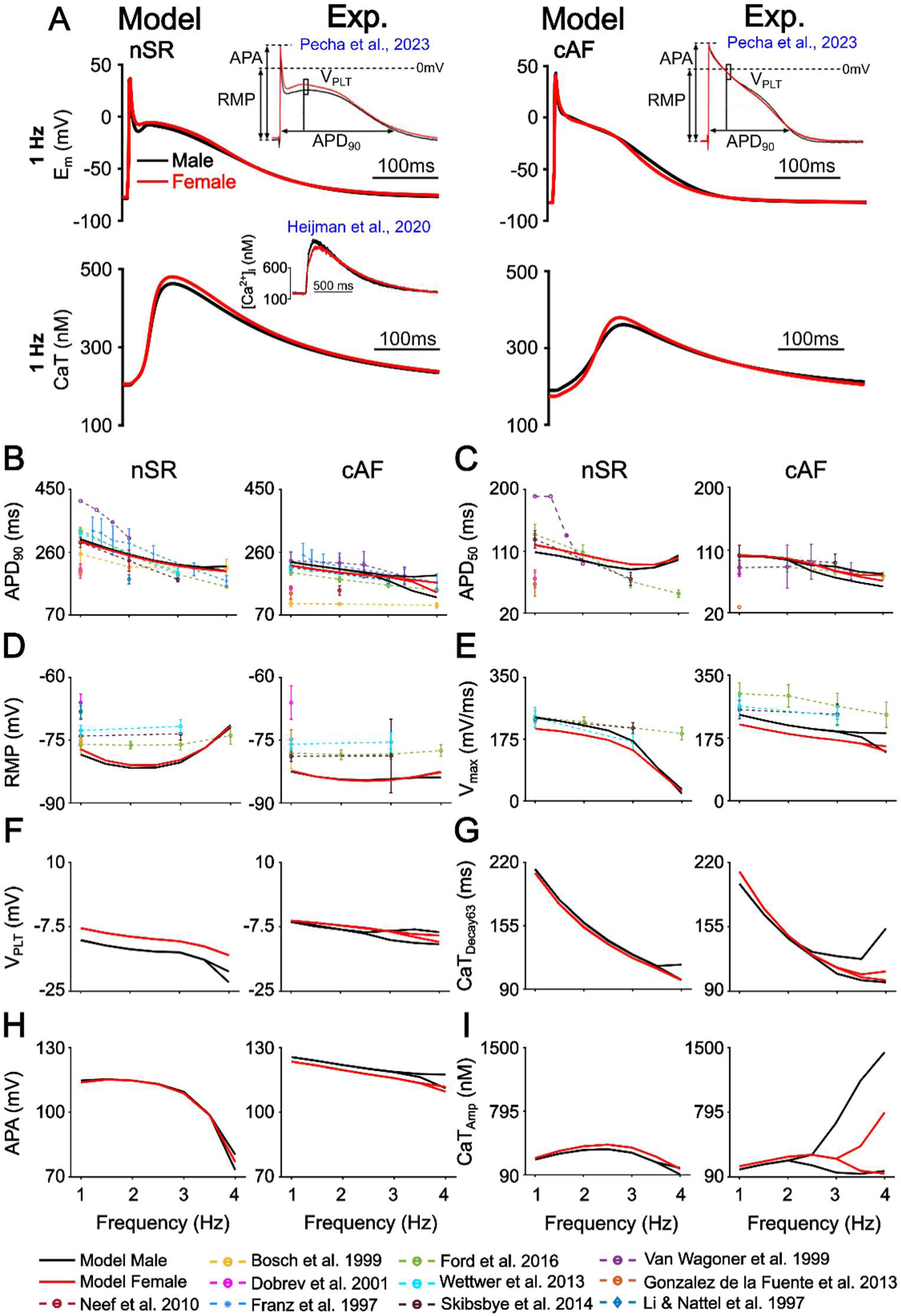
Validation of male and female nSR and female cAF human atrial cardiomyocyte models. *A*) Simulated male and female AP waveforms are compared with sex-specific recordings from human right atrial trabeculae in nSR and cAF (Pecha *et al*., 2023). Simulated male and female CaT traces in nSR are compared with sex-specific recordings from human atrial cardiomyocytes (Heijman *et al*., 2020). *B*) Rate-dependent changes in APD_90_ (AP duration at 90% repolarization), *C*) APD_50_, *D*) resting membrane potential (RMP), *E*) maximum upstroke velocity (V_max_), *F*) plateau voltage (V_PLT_), *G*) calcium transient decay at 63% recovery (CaT_Decay63_), *H*) AP amplitude (APA), *I*) CaT amplitude (CaT_Amp_). Model simulation data collected at steady state from 1 to 4 Hz displayed as solid lines (in 0.5 Hz increments). Data from previous studies in figures *A-D* are displayed as dotted lines (Franz *et al*., 1997; Li and Nattel, 1997; Bosch *et al*., 1999; Van Wagoner *et al*., 1999; Dobrev *et al*., 2001; Neef *et al*., 2010; González de la Fuente *et al*., 2013; Wettwer *et al*., 2013; Skibsbye *et al*., 2014; Ford *et al*., 2016). Note that, when the nSR and cAF models develop alternans two alternating beats are shown.

We next assessed the rate dependence of APD_90_, CV, ERP, and WL in homogeneous 1D strands of nSR and cAF male and female cell models. **Fig. 2*A*** shows an example of the S1-S2 protocol, where the left panel illustrates conduction failure when the S2 interval is shorter than or equal to the ERP, while the right panel demonstrates successful propagation following an S2 stimulus delivered at an interval longer than the ERP. In both nSR and cAF, no sex-specific differences were observed in APD_90_, CV, ERP, or WL. In the cAF condition, both sexes exhibited decreased APD_90_, CV, ERP, and WL compared to their corresponding nSR models, reflecting increased substrate vulnerability to AF. The simulated CV values were consistent with experimental data (Hansson *et al*., 1998; Krul *et al*., 2015), as was the simulated ERP (Skibsbye *et al*., 2014; Ford *et al*., 2016) in both nSR and cAF. Predicted WL closely matched experimental data in cAF (Botteron and Smith, 1996). No notable sex differences were observed in the simulated CV, ERP, and WL.

**Figure 2.**
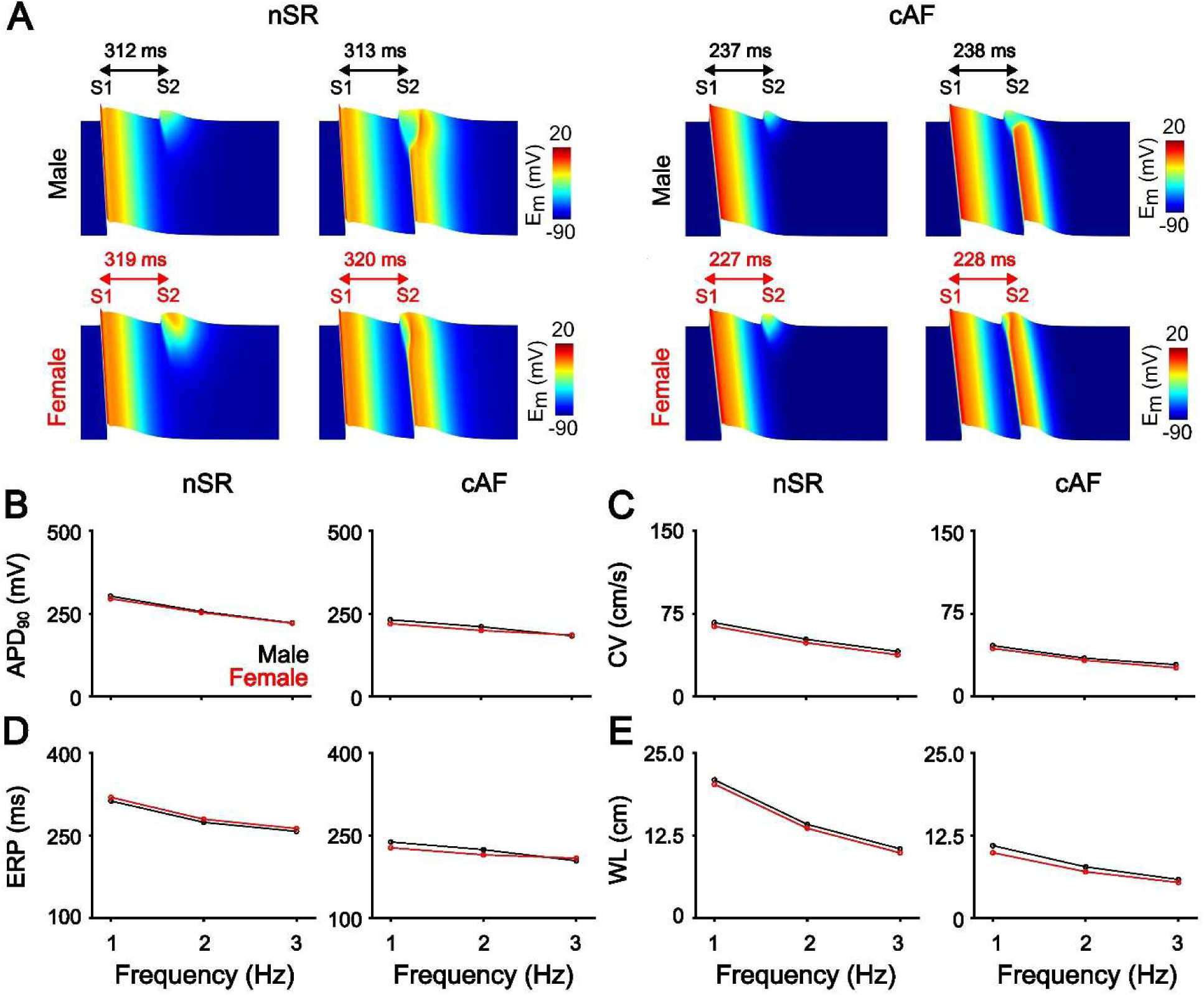
Shorter effective refractory period and wavelength in cAF compared to nSR. *A*) Example of AP propagation in sex-specific nSR and cAF one dimensional strands paced at 1 Hz and measurement of effective refractory period (ERP). Frequency dependence of *B*) AP duration at 90% repolarization (APD_90_), *C*) conduction velocity (CV), *D*) ERP, *E*) and wavelength (WL). In cAF males paced at 3 Hz, AP alternans were observed, with a 17 ms difference in APD_90_ between alternating beats. As a result, the reported values for APD_90_, ERP, CV, and WL represent the average of the two alternating beats.

### Sex-specific parameters contributing to EP differences

We used linear regression analysis (Sobie, 2009) to identify which parameter differences between male and female models in nSR contributed most to the observed variations in key electrophysiological biomarkers (**Fig. 3*A***). The analysis revealed that the longer APD_50_ observed in female vs. male nSR is associated with reduced maximal conductance of the Na^+^ current (G_Na_), ultra-rapid rectifying K^+^ current (G_Kur_), transient outward K^+^ current (G_to_), and increased Na^+^/K^+^ ATPase current maximum velocity (V_NaK_, **Fig. 3*B***). The lower V_max_ in female vs. male nSR correlated with lower G_Na_, G_K1_ and G_Kur_ (**Fig. 3*B***). The more hyperpolarized RMP in female vs. male nSR was linked to reduced G_K1_ and lower CSQ levels (**Fig. 3*B***). The elevated V_PLT_ in female vs. male nSR was associated with decreased G_Kur_, G_to_, and increased V_NaK_ (**Fig. 3*B***).

**Figure 3.**
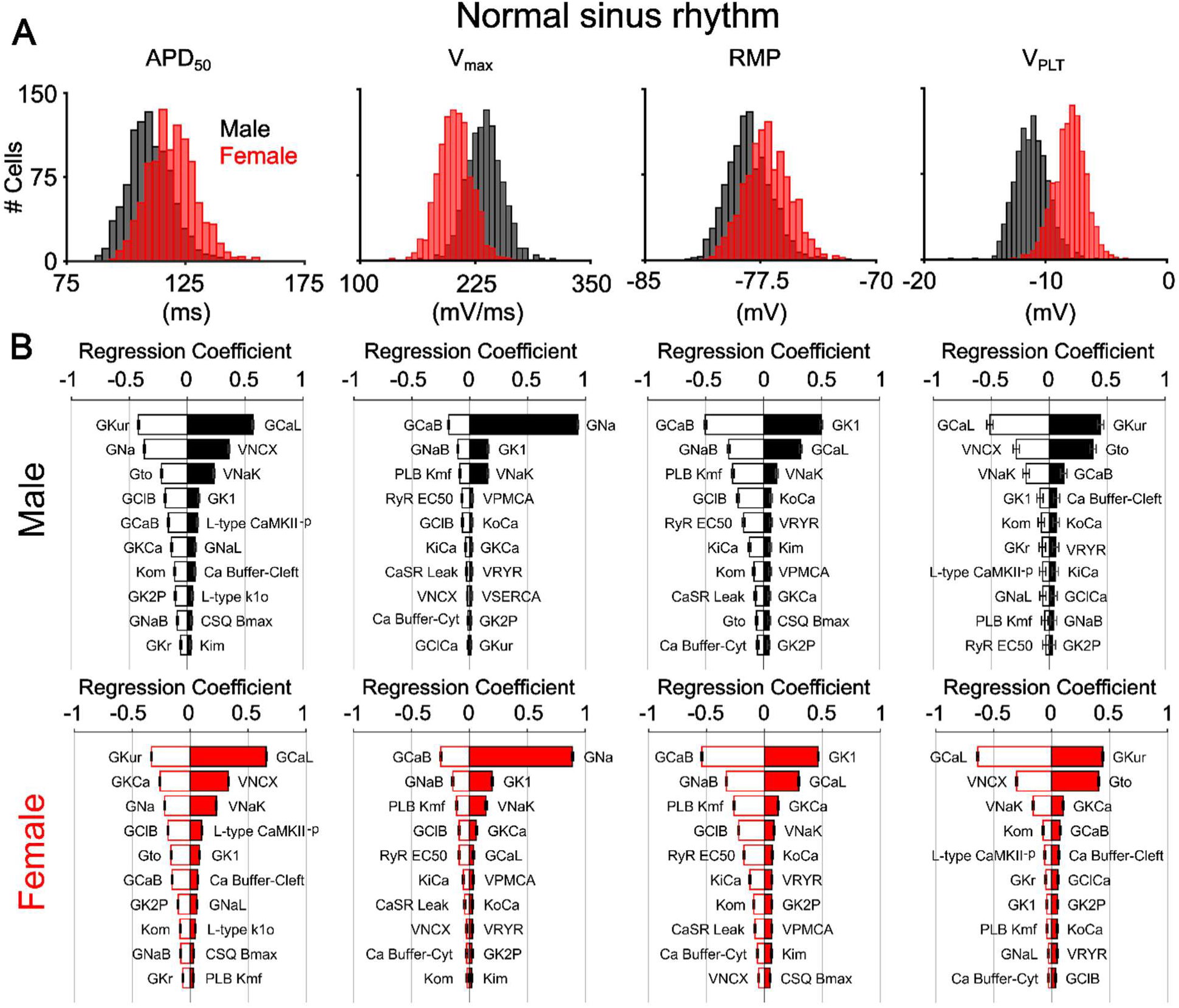
Key parameters associated with sex differences in nSR atrial electrophysiology. *A*) Histograms showing the distribution of nSR male and female AP duration at 50% repolarization (APD_50_), maximum upstroke velocity (V_max_), resting membrane potential (RMP) and plateau voltage (V_PLT_) assessed at 1 Hz pacing in a population of 1,000 models. *B*) Results of linear regression analysis quantifying the sensitivity of each biomarker to changes in the top 20 influential parameters. In all diagrams, a positive coefficient indicates that the increase in the parameter correlates with an increase in the biomarker, whereas a negative coefficient indicates that the increase in the parameter correlates with a decrease in the biomarker. Error bars in the plots represent the 95% CIs, and horizontal ticks were included to indicate CI boundaries.

### Arrhythmia vulnerability in male and female atrial cardiomyocytes under nSR and cAF conditions

We evaluated sex-specific susceptibility to arrhythmogenic alternans and DADs. Alternans, characterized by beat-to-beat oscillations in APD or CaT_Amp_ often precede AF onset (Comtois and Nattel, 2012). In nSR, APD and CaT_Amp_ alternans developed at similar BCLs in males and females (252 ms vs. 230 ms, respectively). In contrast, in cAF, alternans onset occurred at a longer BCL in males than in females (402 ms vs. 305 ms), indicating that males are more vulnerable to alternans in the remodeled state (**Fig. 4*A-B***). At a pacing rate of 300 ms BCL, only cAF males exhibited APD and CaT_Amp_ alternans (**Fig. 4*A***). This finding aligns with clinical observations, primarily in men, showing alternans occur at a higher cycle length in cAF vs. nSR (Narayan *et al*., 2011). To generalize these findings, we determined the APD alternans BCL threshold across our populations of male and female cAF models (**Fig. 4*C***) and applied linear regression analysis to identify the model parameter changes that most strongly correlate with alternans susceptibility (**Fig. 4*D***, Sobie, 2009). Parameters with positive regression coefficients increased the BCL required for alternans onset (i.e., promoted vulnerability), whereas those with negative coefficients were protective. In cAF males, greater alternans vulnerability was associated with reduced G_CaL_, while female protection was associated with an increased activation rate constant for RyR (KoCa) and an increased RyR luminal Ca^2+^ affinity (reduced EC50). Additionally, decreased CSQ expression was associated with reduced alternans susceptibility in cAF females; however, since CSQ was also reduced in cAF males, this alone cannot account for the reduced alternans susceptibility in females. Instead, the combination of reduced CSQ and increased RyR sensitivity to Ca^2+^ likely contributes to the lower alternans propensity observed in cAF females compared to males.

**Figure 4.**
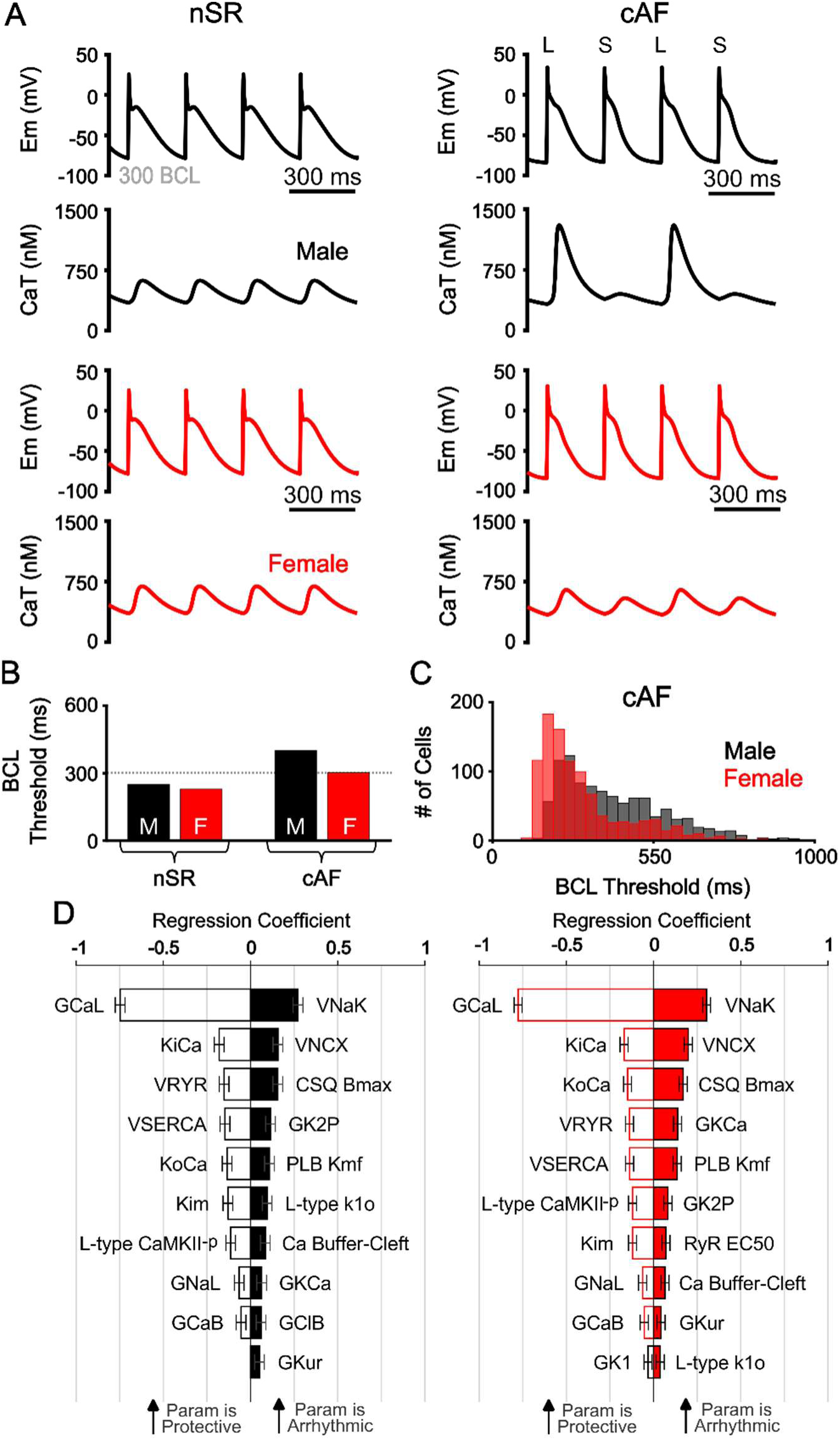
Key parameters associated with vulnerability to APD alternans in male and female cAF. *A*) Time course of simulated male and female nSR and cAF AP and CaT during steady-state pacing at 300 ms basic cycle length (BCL), L - Long AP; S - Short AP. *B*) BCL threshold for alternans (i.e., the longest BCL inducing beat-to-beat APD alternans) in male and female nSR and cAF models. Dashed line indicates 300 ms BCL, which is used in the representative simulations in panel *A*. *C*) In cAF, the BCL threshold for alternans was assessed for 1,000 male and female model variants with randomly varied parameters. Linear regression analysis was used to quantify the sensitivity of the BCL alternans threshold to model parameters, and corresponding regression coefficients are shown in panel *D*. Error bars in the plots represent the 95% CIs, and horizontal ticks were included to indicate CI boundaries.

We next investigated sex-specific differences in the susceptibility to DADs, a known ectopic trigger that contributes to AF initiation. In nSR, the BCL threshold for DAD occurrence was similar between females and males (468 ms vs. 454 ms, respectively). In cAF, however, the BCL threshold was greater in females while remaining unchanged in males (564 ms vs. 453 ms, respectively), indicating a higher vulnerability to DADs in females under cAF conditions (**Fig. 5*B***). As shown in **Fig 5*A***, when paced at 500 ms BCL, cAF females exhibited multiple spontaneous Ca^2+^ release (SCR) events and DADs, whereas males did not. We quantified DAD thresholds in our populations of 1,000 male and 1,000 female cAF models and performed linear regression analysis to identify the parameters most strongly associated with DAD propensity (**Fig. 5*D***, Sobie, 2009). Interestingly, the regression results revealed that many parameters that increased DAD risk were protective against alternans and vice versa. The increased DAD risk in cAF females was associated with enhanced RyR activation rate (KoCa) and increased RyR Ca^2+^ affinity (reduced EC50). In contrast, reduced G_CaL_ in males appeared protective against DADs (**Fig. 5*D***). Although decreased CSQ expression was associated with increased DAD susceptibility in cAF females, CSQ was also decreased in cAF males, who did not show elevated DAD risk. This suggests that reduced CSQ alone is insufficient to explain the sex-specific increase in DADs, but it may contribute to a proarrhythmic substrate in females when combined with other factors, such as increased RyR sensitivity to Ca^2+^.

**Figure 5.**
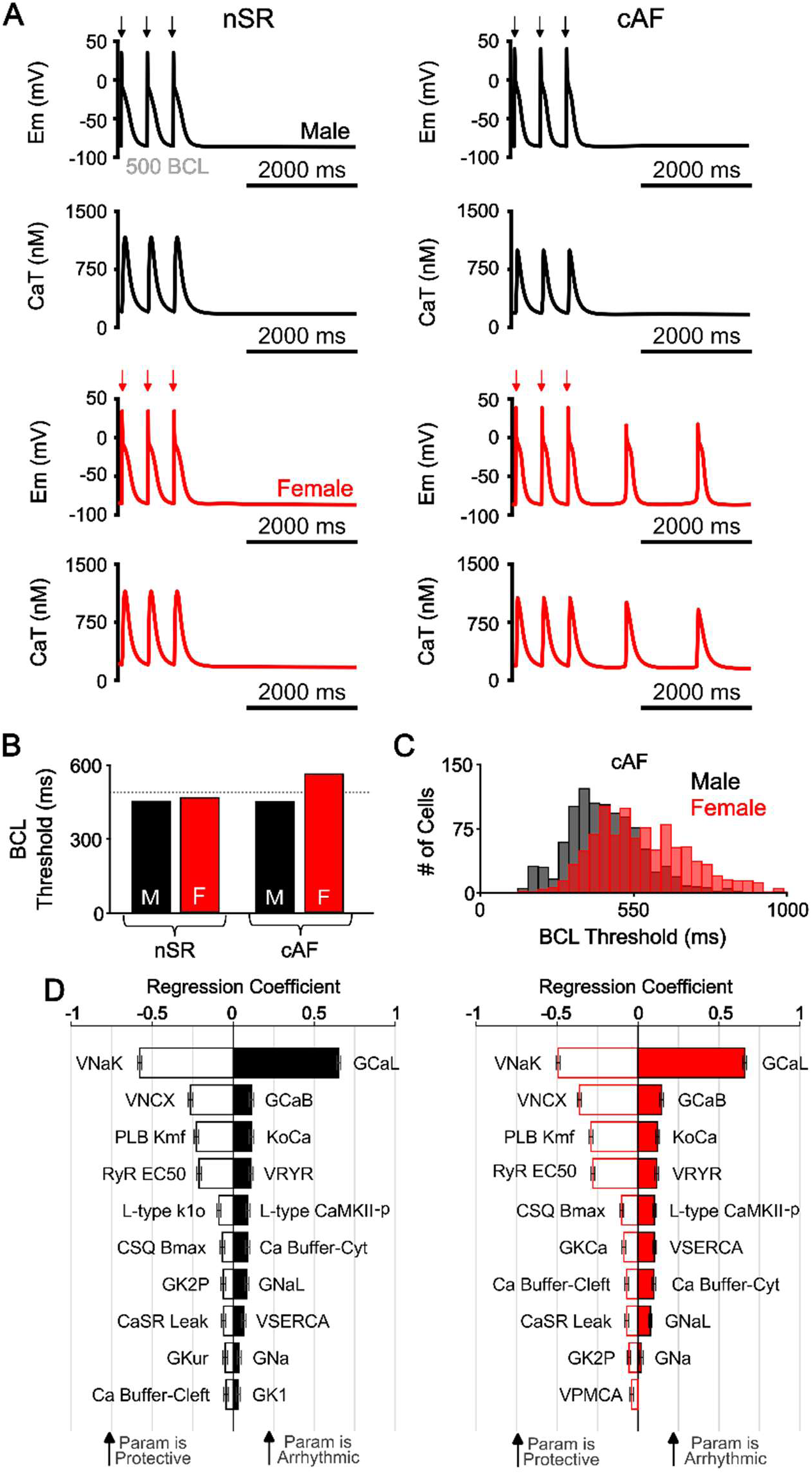
Key parameters associated with DAD propensity in male and female cAF. *A*) Time course of simulated male and female nSR and cAF AP and CaT during steady-state pacing at 500 ms basic cycle length (BCL) and subsequent pause in the presence of Isoproterenol (1 µM). *B*) BCL threshold for DADs (i.e., the longest BCL inducing DADs) in male and female nSR and cAF models. Dashed line indicates 500 ms BCL, which is used in the representative simulations in panel A. *C*) In cAF, the BCL threshold for DADs was assessed for 1,000 male and female model variants with randomly varied parameters. Linear regression analysis was used to calculate regression coefficients quantifying the sensitivity of DAD BCL threshold to model parameters and corresponding regression coefficients are shown in panel *D*. Error bars in the plots represent the 95% CIs, and horizontal ticks were included to indicate CI boundaries.

### Sex-specific treatment strategies for cAF

To better understand sex-specific treatment responses in cAF, we evaluated the effects of several established and proposed therapies, both individually and in combination, building on prior findings that simultaneous inhibition of multiple atrial-predominant ion currents can yield synergistic antiarrhythmic effects (Ni *et al*., 2020). We investigated the antiarrhythmic effects of inhibiting key atrial K^+^ currents (25% reduction in G_SK_ and G_K2P_), both alone and in combination with flecainide’s effects on I_Kr_, I_Na_, I_CaL_, and RyR (**Fig. 6*A***). Our therapeutic goal was to reverse key cAF-induced remodeling features, specifically, the shortened APD_90_ and reduced CaT_Amp_, bringing them closer to sex-specific nSR levels, while also reducing DAD propensity in females (i.e., decreasing the DAD BCL threshold) and alternans vulnerability in males (i.e., decreasing the alternans BCL threshold). Dotted grey lines in **Fig. 6*B-E*** represent complete restoration of each biomarker to nSR values. In cAF males, a wider range of drug combinations proved effective. The most effective intervention was flecainide at 1x or 2x the EFTPC combined with G_SK_ and G_K2P_ inhibition, which substantially increased APD_90_, slightly increased CaT_Amp_, modestly lowered the BCL threshold for alternans (especially at 1x EFTPC), and had minimal impact on DAD development (**Fig. 6*B-E***). In cAF females, only the combination of flecainide at 1x EFTPC with G_SK_ and G_K2P_ inhibition produced beneficial effects, including a substantial increase in APD_90_, a modest increase in CaT_Amp_, no change in the alternans BCL threshold, and a slight reduction in DAD propensity (**Fig. 6*B-E***).

**Figure 6.**
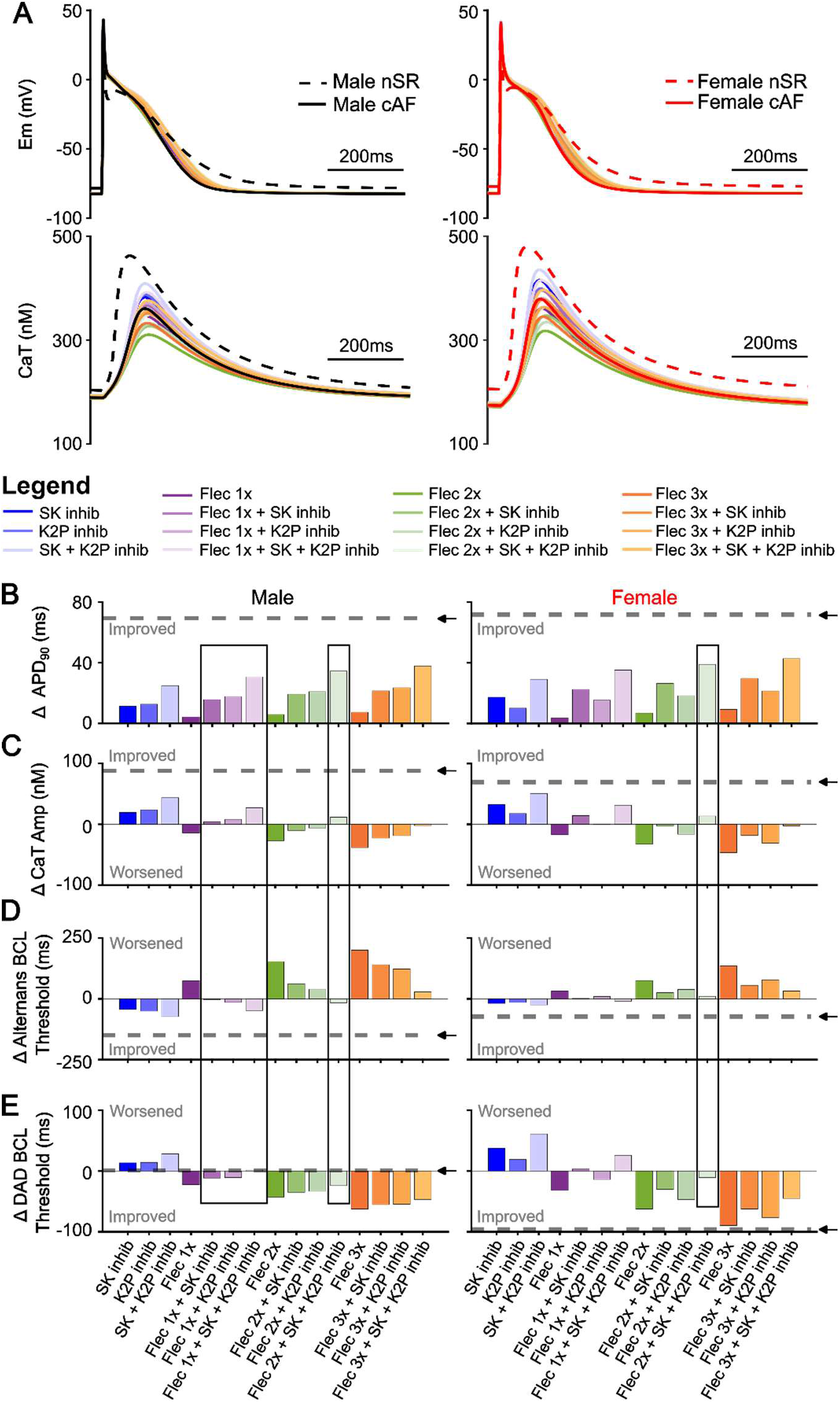
Sex-specific impact of SK and K2P channel inhibition and flecainide on human atrial electrophysiology, Ca^2+^ handling, and arrhythmia vulnerability. *A*) Male and female AP and CaT traces at 1 Hz in nSR, cAF, and cAF following various drug regiments, including 25% inhibition of SK (G_SK_), K2P (G_K2P_), G_SK_ + G_K2P_, and flecainide (Flec) at 1x, 2x, and 3x effective free therapeutic plasma concentrations, both alone and in combination with G_SK_ and/or G_K2P_ inhibition. For each drug condition, we quantified the effect on AP duration at 90% repolarization (APD_90_), CaT amplitude (CaT_Amp_), and basic cycle length (BCL) thresholds for the onset of alternans and delayed afterdepolarizations (DADs). Panels *B-E* report the difference of each biomarker between the drug condition and baseline cAF. Dotted grey lines represent the difference between nSR and cAF for each sex-specific biomarker. Black boxes indicate effective therapeutics in male and female cAF, whereby there is prolongation of APD_90_, increase in CaT_Amp_, and reduced arrhythmia vulnerability.

## Discussion

A deeper mechanistic understanding of sex-specific mechanisms in AF is essential to inform the development of more effective, individualized treatment strategies. In this study we developed and validated novel sex-specific human cardiomyocyte models recapitulating differences in atrial EP and Ca^2+^ cycling under nSR and cAF conditions. Our simulations revealed distinct sex-specific patterns of arrhythmia vulnerability and treatment response in cAF. Simulated drug treatments revealed a broader range of effective interventions in males, particularly combinations involving flecainide and K^+^-channel blockade, whereas females exhibited limited therapeutic benefit. These results suggest that the mechanisms driving arrhythmia differ by sex and underscore the importance of developing sex-specific strategies for AF management.

### Molecular Determinants of Sex Differences in Electrophysiological Biomarkers and Arrhythmogenic Substrate

Sex differences in atrial AP features have been reported in humans (Ravens, 2018; Pecha *et al*., 2023) at 1 Hz, yet the mechanisms underlying these differences remain poorly defined. While our previous work focused on subcellular Ca^2+^ cycling under AP clamp (Zhang *et al*., 2024), the present study provides new insights by identifying key model parameters associated with these AP differences. In nSR at 1 Hz, females exhibited longer APD_50_, higher V_PLT_, reduced V_max_, and more depolarized RMP primarily driven by lower G_Na_, G_Kur_, G_K1_, G_to_, and higher V_NaK_ (**Fig. 3**). These differences diminished at higher pacing rates and were largely attenuated in cAF. Although atrial remodeling in cAF vs. nSR is similar in males and females, with both showing shortened APD, reduced ERP, CV and WL, other aspects of the arrhythmogenic substrate diverge between sexes. In cAF, males were more prone to APD and CaT_Amp_ alternans, while females showed greater susceptibility to DADs. This suggests that although cAF-induced electrical remodeling reduces observable sex differences in AP features, underlying sex-specific changes may continue to drive differences in arrhythmia susceptibility. Sex-specific differences persisted in the underlying determinants of arrhythmogenic susceptibility, with alternans in males being associated with reduced G_CaL_ and increased DAD propensity in females correlating with enhanced RyR sensitivity to Ca^2+^ (**Figs. 4-5**). The latter findings confirm our previous observations (Zhang *et al*., 2024) that RyR phosphorylation and I_CaL_ are key drivers of SCR. Our multivariable analysis similarly identified RyR sensitivity and G_CaL_ as primary determinants of DADs, along with decreasing CSQ or PLB expression. Interestingly, however, decreasing PLB or CSQ, or increasing RyR phosphorylation or I_CaL_ activity increased beat-to-beat variability of the CaT under AP clamp in our previous study (Zhang *et al*., 2024). In the present study, when Ca^2+^ and membrane voltage are coupled, the same perturbations are associated with a decreased alternans BCL threshold, underscoring how AP-Ca^2+^ interactions alter alternans vulnerability.

While these results reflect our sex-specific parameterization, which may be influenced by confounding factors (Herraiz-Martínez *et al*., 2022) as discussed in the Methods, they illustrate that the magnitude of G_CaL_ is a key determinant of both alternans susceptibility and DAD propensity, regardless of sex, and that shifting this single parameter alters arrhythmia vulnerability in opposing directions. Specifically, G_CaL_ reduction decreases DAD risk but promotes alternans, whereas elevated G_CaL_ reduces the susceptibility to alternans but promotes DADs by facilitating Ca²⁺ overload. Similarly, increased RyR activation promotes DADs but can mitigate alternans (and vice versa). These antagonistic relationships suggest that a given therapeutic intervention may have protective or pro-arrhythmic effects depending on the dominant arrhythmia mechanism and the biological sex. In this context, our results suggest that if males with cAF do experience greater I_CaL_ reduction, this could help explain their greater alternans vulnerability and lower DAD risk. Conversely, if females exhibit elevated RyR sensitivity or phosphorylation, this could increase DAD risk. However, definitive conclusions require more robust, sex-stratified experimental studies to validate the presence and magnitude of these remodeling differences in human atrial tissue.

### Sex-Specific Pharmacological Responses to Antiarrhythmic Therapy

Emerging therapeutic strategies for AF have increasingly focused on atrial-predominant targets, including inhibition of SK channels (Heijman *et al*., 2023; Holst *et al*., 2024) and K2P channels (Schmidt *et al*., 2014). These strategies hold promise for reducing ventricular side effects, but their efficacy across sexes remains poorly understood. This knowledge gap is particularly concerning given accumulating evidence of sex differences in arrhythmia mechanisms, which limits our understanding of sex-specific responses to antiarrhythmic therapies.

We previously investigated mechanisms underlying increased spontaneous Ca²⁺ release in females using AP clamp simulations (Zhang *et al*., 2024), but could not capture sex-dependent differences in AP morphology or assess the effects of therapeutic interventions on arrhythmia mechanisms such as DADs, alternans, or conduction. Here, we used our validated male and female nSR and cAF models to investigate the effects of flecainide, a clinically used antiarrhythmic agent, alone and in combination with SK and K2P channel inhibition. These interventions were selected based on prior evidence suggesting that multi-channel targeting can yield synergistic antiarrhythmic effects (Ni *et al*., 2017; Dasí *et al*., 2024). Our simulations were designed to evaluate the ability of these interventions to reverse cAF-induced remodeling and mitigate sex-specific arrhythmias. The results revealed marked differences in treatment efficacy. In males, several drug combinations, particularly flecainide with SK and K2P inhibition, were effective in prolonging APD_90_, increasing CaT_Amp_, and lowering the threshold for alternans. In contrast, the same combinations had only modest effects in females, producing limited improvements in APD_90_ and CaT_Amp_, and minimal reduction in DAD susceptibility. These limited effects suggest that current strategies may not adequately address the Ca^2+^-driven arrhythmia mechanisms more prominent in females, particularly those linked to enhanced RyR sensitivity to Ca^2+^. The distinct arrhythmogenic mechanisms identified in males and females may also have implications for non-pharmacological therapies such as catheter ablation. For instance, alternans-driven reentry, more prevalent in males, may respond better to strategies that target conduction pathways and reentrant circuits. In contrast, DAD-mediated triggers, which predominate in females, may require more extensive substrate modification or adjunctive therapies to suppress focal activity. These differences could partly explain the lower success rates and higher recurrence observed in women following ablation (Patel *et al*., 2010; Watanabe *et al*., 2022).

Together, these findings reinforce that males and females not only differ in their underlying arrhythmogenic substrates but also in their responses to treatment. To develop effective and equitable AF treatments it is essential to account for these sex-specific differences and to address the persistent underrepresentation of women in both basic and clinical cardiovascular research, which continues to limit our understanding of female-specific responses to antiarrhythmic interventions (Tobb, Kocher and Bullock-Palmer, 2022; Matthews *et al*., 2024).

### Limitations and Future Work

Several limitations should be considered when interpreting the findings of this study. First, the sex-specific models were primarily built on data from human atrial tissue obtained from patients undergoing cardiac surgery. While the individuals in nSR did not present with AF, their atrial tissue may still exhibit pathological alterations, which could influence baseline electrophysiological and Ca^2+^ handling properties. Similarly, the cAF remodeling reflects both chronic AF remodeling and alterations related to comorbidities and/or ablation procedure. Future studies with larger sample sizes and sex-stratified designs are needed.

Second, integration of sex-specific features into multi-scale models spanning subcellular to cell to tissue level is necessary to advance our understanding of arrhythmia vulnerability in males and females. Prior studies have documented morphological differences between male and female cardiomyocytes, including cell size and t-tubule organization (Yue *et al*., 2017; Thibault *et al*., 2022), which we have investigated in prior work (Zhang *et al*., 2024). Additionally, tissue-scale sex differences, including fibrosis that is more pronounced in female cAF atria (Akoum *et al*., 2018), may critically shape arrhythmia vulnerability and incorporating these sex-specific features will be essential for capturing the spatial and structural substrates that modulate AF onset and maintenance in a sex-dependent manner and informing sex-specific therapeutic strategies. Moreover, atrial repolarization exhibits considerable spatial heterogeneity under both physiological and pathological conditions, and remodeling is unlikely to be uniform across atrial regions. These spatial differences could amplify regional disparities in APD, Ca^2+^ handling, and conduction, thereby creating localized zones of heightened arrhythmogenicity.

Third, while our findings support the benefit of multi-target interventions, further work is needed to optimize sex-specific drug combinations and assess their efficacy in tissue-level models. Future studies should also investigate the role of sex hormones in modulating atrial structure, ion channel expression, and EP. For example, testosterone deficiency has been strongly associated with increased AF risk in men (Magnani *et al*., 2014). The effects of estrogen and progesterone on AF susceptibility in women remains poorly understood (Perez *et al*., 2012; Tsai *et al*., 2016; Wong *et al*., 2017; Lee *et al*., 2021), but emerging evidence suggests that estrogen may exert protective effects by limiting atrial fibrosis (Regitz-Zagrosek, 2020), which is a key contributor to arrhythmia maintenance. Inflammation, another driver of structural remodeling in AF, is also influenced by sex and sex hormones, with estrogen being shown to exert anti-inflammatory effects that may further mitigate fibrotic remodeling (Mclarty *et al*., 2013). In parallel, multiple studies have shown that sex hormones modulate ion channel function and atrial EP (Nakajima *et al*., 1999; Tsuneda *et al*., 2009; Papp *et al*., 2017; Giammarino *et al*., 2025). Incorporating hormonal regulation of these processes into sex-specific models may provide critical insights into the endocrine contributions to AF pathophysiology and help identify novel targets for personalized therapeutic intervention.

Finally, while not explicitly modeled in this study, age and comorbidities such as hypertension and diabetes are known to alter cardiac EP and may interact with sex-specific remodeling in AF. Future work should incorporate these factors to better reflect patient diversity and improve clinical relevance.

### Conclusions

This study demonstrates that sex-specific aspects of cAF-associated ionic remodeling may predispose males and females to distinct arrhythmia mechanisms. Although our model assumptions are based on limited experimental data and require further validation in larger, sex-stratified patient cohorts, the simulations illustrate how such differences can shape arrhythmia vulnerability and influence treatment response. Notably, I_CaL_ remodeling emerged as a key determinant of arrhythmia susceptibility, with its reduction influencing the balance between alternans and DADs. Given known inter-individual variability, the extent and spatial pattern of I_CaL_ downregulation may critically define patients’ arrhythmic risk profile. Pharmacological simulations revealed greater treatment efficacy in males, highlighting the need for sex-specific models to guide precision therapies for AF. Together, these results emphasize the importance of incorporating sex as a biological variable in AF modeling, clinical trial and treatment design to accurately assess antiarrhythmic drug efficacy and improve precision medicine strategies for both men and women.

## Additional information section

## Data Availability Statement

Our codes, as well as their comprehensive description, are freely available at https://github.com/drgrandilab/.

## Author contributions

All authors contributed to the conception and design of the work and participated in drafting the work or critically revising it for important intellectual content. All authors have read and approved the final version of the manuscript. In addition, all authors agree to be accountable for all aspects of the work in ensuring that questions related to the accuracy or integrity of any part of the work are appropriately investigated and resolved. All persons designated as authors qualify for authorship, and all those that qualify for authorship are listed.

## Funding

American Heart Association Predoctoral Fellowship 24PRE1183427 (NTH)

American Heart Association Career Development Award 24CDA1258695 (HN)

American Heart Association Postdoctoral Fellowship 20POST35120462 (HN)

American Heart Association Career Development Award 24CDA1269250 (CERS)

NIA Grant R03AG086695 (CERS, EG)

NHLBI Grant R01HL176651 (EG, DD, SM)

NHLBI Grants R01HL131517, R01HL141214, and P01HL141084 (EG)

NHLBI Grant R01HL170521 (EG, HN)

NHLBI Grants R00HL138160, R01HL171057, and R01HL171586 (SM)

NHLBI Grants R01HL136389, R01HL163277, R01HL160992, R01HL165704, and R01HL164838 (DD)

Deutsche Forschungsgemeinschaft Research Training Group 2989 (DD)

European Union large-scale network grant No. 965286 (MAESTRIA, DD)

National Institutes of Health Stimulating Peripheral Activity to Relieve Conditions Grant 1OT2OD026580-01 (EG)

## References

Akoum, N. et al. (2018) ‘Age and sex differences in atrial fibrosis among patients with atrial fibrillation’, Europace: European Pacing, Arrhythmias, and Cardiac Electrophysiology: Journal of the Working Groups on Cardiac Pacing, Arrhythmias, and Cardiac Cellular Electrophysiology of the European Society of Cardiology, 20(7), pp. 1086–1092. Available at: 10.1093/europace/eux260.

Ambrosi, C.M. et al. (2013) ‘Gender Differences in Electrophysiological Gene Expression in Failing and Non-Failing Human Hearts’, PLoS ONE, 8(1), p. e54635. Available at: 10.1371/journal.pone.0054635.

Ball, J. et al. (2013) ‘Women versus men with chronic atrial fibrillation: insights from the Standard versus Atrial Fibrillation spEcific managemenT studY (SAFETY)’, PloS One, 8(5), p. e65795. Available at: 10.1371/journal.pone.0065795.

Bosch, R.F. et al. (1999) ‘Ionic mechanisms of electrical remodeling in human atrial fibrillation’, Cardiovascular Research, 44(1), pp. 121–131. Available at: 10.1016/S0008-6363(99)00178-9.

Botteron, G.W. and Smith, J.M. (1996) ‘Quantitative Assessment of the Spatial Organization of Atrial Fibrillation in the Intact Human Heart’, Circulation, 93(3), pp. 513–518. Available at: 10.1161/01.CIR.93.3.513.

Brundel, B.J.J.M. et al. (2001) ‘Ion Channel Remodeling Is Related to Intraoperative Atrial Effective Refractory Periods in Patients With Paroxysmal and Persistent Atrial Fibrillation’, Circulation, 103(5), pp. 684–690. Available at: 10.1161/01.CIR.103.5.684.

Caballero, R. et al. (2010) ‘In Humans, Chronic Atrial Fibrillation Decreases the Transient Outward Current and Ultrarapid Component of the Delayed Rectifier Current Differentially on Each Atria and Increases the Slow Component of the Delayed Rectifier Current in Both’, Journal of the American College of Cardiology, 55(21), pp. 2346–2354. Available at: 10.1016/j.jacc.2010.02.028.

Comtois, P. and Nat℡, S. (2012) ‘Atrial Repolarization Alternans as a Path to Atrial Fibrillation’, Journal of Cardiovascular Electrophysiology, 23(9), pp. 1013–1015. Available at: 10.1111/j.1540-8167.2012.02391.x.

Dasí, A. et al. (2024) ‘Prospective in silico trials identify combined SK and K2P channel block as an effective strategy for atrial fibrillation cardioversion’, The Journal of Physiology [Preprint]. Available at: 10.1113/JP287124.

Dobrev, D. et al. (2001) ‘Molecular Basis of Downregulation of G-Protein–Coupled Inward Rectifying K+ Current (IK,ACh) in Chronic Human Atrial Fibrillation’, Circulation, 104(21), pp. 2551–2557. Available at: 10.1161/hc4601.099466.

Dobrev, D. and Ravens, U. (2003) ‘Remodeling of cardiomyocyte ion channels in human atrial fibrillation’, Basic Research in Cardiology, 98(3), pp. 137–148. Available at: 10.1007/s00395-003-0409-8.

El-Armouche, A. et al. (2006) ‘Molecular determinants of altered Ca2+ handling in human chronic atrial fibrillation’, Circulation, 114(7), pp. 670–680. Available at: 10.1161/CIRCULATIONAHA.106.636845.

Essebag, V. et al. (2007) ‘Sex Differences in the Relationship Between Amiodarone Use and the Need for Permanent Pacing in Patients With Atrial Fibrillation’, Archives of Internal Medicine, 167(15), pp. 1648–1653. Available at: 10.1001/archinte.167.15.1648.

Ford, J. et al. (2016) ‘The positive frequency-dependent electrophysiological effects of the IKur inhibitor XEN-D0103 are desirable for the treatment of atrial fibrillation’, Heart Rhythm, 13(2), pp. 555–564. Available at: 10.1016/j.hrthm.2015.10.003.

Franz, M.R. et al. (1997) ‘Electrical Remodeling of the Human Atrium: Similar Effects in Patients With Chronic Atrial Fibrillation and Atrial Flutter 1’, Journal of the American College of Cardiology, 30(7), pp. 1785–1792. Available at: 10.1016/S0735-1097(97)00385-9.

Gaborit, N. et al. (2010) ‘Gender-related differences in ion-channel and transporter subunit expression in non-diseased human hearts’, Journal of Molecular and Cellular Cardiology, 49(4), pp. 639–646. Available at: 10.1016/j.yjmcc.2010.06.005.

Giammarino, L. et al. (2025) ‘Sex and sex hormonal regulation of the atrial inward rectifier potassium current (IK1): insights into potential pro-arrhythmic mechanisms’, *Cardiovascular Research*, p. cvaf074. Available at: 10.1093/cvr/cvaf074.

González de la Fuente, M., et al. (2013) ‘Chronic atrial fibrillation up-regulates β1-Adrenoceptors affecting repolarizing currents and action potential duration’, Cardiovascular Research, 97(2), pp. 379–388. Available at: 10.1093/cvr/cvs313.

Grandi, E. et al. (2011) ‘Human atrial action potential and Ca2+ model: sinus rhythm and chronic atrial fibrillation’, Circulation Research, 109(9), pp. 1055–1066. Available at: 10.1161/CIRCRESAHA.111.253955.

GTEx Consortium (2020) ‘The GTEx Consortium atlas of genetic regulatory effects across human tissues’, *Science (New York*, N.Y.), 369(6509), pp. 1318–1330. Available at: 10.1126/science.aaz1776.

Hansson, A. et al. (1998) ‘Right atrial free wall conduction velocity and degree of anisotropy in patients with stable sinus rhythm studied during open heart surgery’, European Heart Journal, 19(2), pp. 293–300. Available at: 10.1053/euhj.1997.0742.

Heijman, J., Muna, A.P., Veleva, T., Molina, C.E., Sutanto, H., Tekook, M., Wang, Q., Abu-Taha, I.H., Gorka, M., Künzel, S., El-Armouche, A., Reichenspurner, H., Kamler, M., Nikolaev, V., Ravens, U., Li, N., Nattel, S., Wehrens, Xander H. T., et al. (2020) ‘Atrial Myocyte NLRP3/CaMKII Nexus Forms a Substrate for Postoperative Atrial Fibrillation’, Circulation Research, 127(8), pp. 1036–1055. Available at: 10.1161/CIRCRESAHA.120.316710.

Heijman, J., Muna, A.P., Veleva, T., Molina, C.E., Sutanto, H., Tekook, M., Wang, Q., Abu-Taha, I.H., Gorka, M., Künzel, S., El-Armouche, A., Reichenspurner, H., Kamler, M., Nikolaev, V., Ravens, U., Li, N., Nattel, S., Wehrens, Xander H.T., et al. (2020) ‘Atrial Myocyte NLRP3/CaMKII Nexus Forms a Substrate for Post-Operative Atrial Fibrillation’, Circulation research, 127(8), pp. 1036–1055. Available at: 10.1161/CIRCRESAHA.120.316710.

Heijman, J. et al. (2023) ‘Enhanced Ca2+-Dependent SK-Channel Gating and Membrane Trafficking in Human Atrial Fibrillation’, Circulation Research, 132(9), pp. e116–e133. Available at: 10.1161/CIRCRESAHA.122.321858.

Herraiz-Martínez, A. et al. (2022) ‘Influence of sex on intracellular calcium homoeostasis in patients with atrial fibrillation’, Cardiovascular Research, 118(4), pp. 1033–1045. Available at: 10.1093/cvr/cvab127.

Herrera, N.T. et al. (2023) ‘Dual effects of the small-conductance Ca2+-activated K+ current on human atrial electrophysiology and Ca2+-driven arrhythmogenesis: an in silico study’. bioRxiv, p. 2023.06.16.545367. Available at: 10.1101/2023.06.16.545367.

Holst, A.G. et al. (2024) ‘Inhibition of the KCa2 potassium channel in atrial fibrillation: a randomized phase 2 trial’, Nature Medicine, 30(1), pp. 106–111. Available at: 10.1038/s41591-023-02679-9.

Iseppe, A.F. et al. (2021) ‘Sex-specific classification of drug-induced Torsade de Pointes susceptibility using cardiac simulations and machine learning’, Clinical pharmacology and therapeutics, 110(2), pp. 380–391. Available at: 10.1002/cpt.2240.

Khaing, E. et al. (2025) ‘Representation of Women in Atrial Fibrillation Ablation Randomized Controlled Trials: Systematic Review’, Journal of the American Heart Association, 14(2), p. e035181. Available at: 10.1161/JAHA.124.035181.

Ko, D. et al. (2016) ‘Atrial fibrillation in women: epidemiology, pathophysiology, presentation, and prognosis’, Nature reviews. Cardiology, 13(6), pp. 321–332. Available at: 10.1038/nrcardio.2016.45.

Kornej, J. et al. (2020) ‘Epidemiology of Atrial Fibrillation in the 21st Century’, Circulation Research, 127(1), pp. 4–20. Available at: 10.1161/CIRCRESAHA.120.316340.

Krul, S.P.J. et al. (2015) ‘Atrial fibrosis and conduction slowing in the left atrial appendage of patients undergoing thoracoscopic surgical pulmonary vein isolation for atrial fibrillation’, Circulation. Arrhythmia and Electrophysiology, 8(2), pp. 288–295. Available at: 10.1161/CIRCEP.114.001752.

Lai, L.-P. et al. (2000) ‘Changes in the mRNA Levels of Delayed Rectifier Potassium Channels in Human Atrial Fibrillation’, Cardiology, 92(4), pp. 248–255. Available at: 10.1159/000006982.

Lee, J. et al. (2021) ‘Clinical Impact of Hormone Replacement Therapy on Atrial Fibrillation in Postmenopausal Women: A Nationwide Cohort Study’, Journal of Clinical Medicine, 10(23), p. 5497. Available at: 10.3390/jcm10235497.

Li, G.R. and Nattel, S. (1997) ‘Properties of human atrial ICa at physiological temperatures and relevance to action potential’, American Journal of Physiology-Heart and Circulatory Physiology, 272(1), pp. H227–H235. Available at: 10.1152/ajpheart.1997.272.1.H227.

Magnani, J.W. et al. (2014) ‘Association of Sex Hormones, Aging and Atrial Fibrillation in Men: The Framingham Heart Study’, Circulation. Arrhythmia and electrophysiology, 7(2), pp. 307–312. Available at: 10.1161/CIRCEP.113.001322.

Matthews, S. et al. (2024) ‘Factors affecting women’s participation in cardiovascular research: a scoping review’, European Journal of Cardiovascular Nursing, 23(2), pp. 107–114. Available at: 10.1093/eurjcn/zvad048.

Mclarty, J.L. et al. (2013) ‘Estrogen modulates the influence of cardiac inflammatory cells on function of cardiac fibroblasts’, Journal of Inflammation Research, 6, pp. 99–108. Available at: 10.2147/JIR.S48422.

Morotti, S. and Grandi, E. (2024) ‘Population-Based Computational Approaches to Investigate Cardiac Arrhythmia Risk’, in T. Jue (ed.) Molecular and Computational Modeling of Cardiac Function. Cham: Springer Nature Switzerland, pp. 181–198. Available at: 10.1007/978-3-031-73730-5_5.

Muzzey, M. et al. (2020) ‘Flecainide is well-tolerated and effective in patient with atrial fibrillation at 12 months: a retrospective study’, Therapeutic Advances in Cardiovascular Disease, 14, p. 1753944720926824. Available at: 10.1177/1753944720926824.

Nakajima, T. et al. (1999) ‘Antiarrhythmic effect and its underlying ionic mechanism of 17β-estradiol in cardiac myocytes’, British Journal of Pharmacology, 127(2), pp. 429–440. Available at: 10.1038/sj.bjp.0702576.

Narayan, S.M. et al. (2011a) ‘Repolarization alternans reveals vulnerability to human atrial fibrillation’, Circulation, 123(25), pp. 2922–2930. Available at: 10.1161/CIRCULATIONAHA.110.977827.

Narayan, S.M. et al. (2011b) ‘Repolarization Alternans Reveals Vulnerability to Human Atrial Fibrillation’, Circulation, 123(25), pp. 2922–2930. Available at: 10.1161/CIRCULATIONAHA.110.977827.

Neef, S. et al. (2010) ‘CaMKII-Dependent Diastolic SR Ca2+ Leak and Elevated Diastolic Ca2+ Levels in Right Atrial Myocardium of Patients With Atrial Fibrillation’, Circulation Research, 106(6), pp. 1134–1144. Available at: 10.1161/CIRCRESAHA.109.203836.

Nelder, J.A. and Mead, R. (1965) ‘A Simplex Method for Function Minimization’, The Computer Journal, 7(4), pp. 308–313. Available at: 10.1093/comjnl/7.4.308.

Ni, H. et al. (2017) ‘Synergistic Anti-arrhythmic Effects in Human Atria with Combined Use of Sodium Blockers and Acacetin’, Frontiers in Physiology, 8, p. 946. Available at: 10.3389/fphys.2017.00946.

Ni, H. et al. (2020) ‘Populations of in silico myocytes and tissues reveal synergy of multiatrial-predominant K ^+^-current block in atrial fibrillation’, *British Journal of Pharmacology*, p. bph.15198. Available at: 10.1111/bph.15198.

Ni, H. et al. (2023) ‘Integrative human atrial modelling unravels interactive protein kinase A and Ca2+/calmodulin-dependent protein kinase II signalling as key determinants of atrial arrhythmogenesis’, Cardiovascular Research, 119(13), pp. 2294–2311. Available at: 10.1093/cvr/cvad118.

Odening, K.E. et al. (2019) ‘Mechanisms of sex differences in atrial fibrillation: role of hormones and differences in electrophysiology, structure, function, and remodelling’, EP Europace, 21(3), pp. 366–376. Available at: 10.1093/europace/euy215.

Papp, R. et al. (2017) ‘Genomic upregulation of cardiac Cav1.2α and NCX1 by estrogen in women’, Biology of Sex Differences, 8(1), p. 26. Available at: 10.1186/s13293-017-0148-4.

Patel, D. et al. (2010) ‘Outcomes and complications of catheter ablation for atrial fibrillation in females’, Heart Rhythm, 7(2), pp. 167–172. Available at: 10.1016/j.hrthm.2009.10.025.

Pecha, S. et al. (2023) ‘Resting membrane potential is less negative in trabeculae from right atrial appendages of women, but action potential duration does not shorten with age’, Journal of Molecular and Cellular Cardiology, 176, pp. 1–10. Available at: 10.1016/j.yjmcc.2023.01.006.

Perez, M.V. et al. (2012) ‘Effects of postmenopausal hormone therapy on incident atrial fibrillation: the Women’s Health Initiative randomized controlled trials’, Circulation. Arrhythmia and Electrophysiology, 5(6), pp. 1108–1116. Available at: 10.1161/CIRCEP.112.972224.

Ravens, U. (2018) ‘Sex differences in cardiac electrophysiology’, Canadian Journal of Physiology and Pharmacology, 96(10), pp. 985–990. Available at: 10.1139/cjpp-2018-0179.

Regitz-Zagrosek, V. (2020) ‘Sex and Gender Differences in Heart Failure’, International Journal of Heart Failure, 2(3), pp. 157–181. Available at: 10.36628/ijhf.2020.0004.

Rienstra, M. et al. (2012) ‘Symptoms and Functional Status of Patients with Atrial Fibrillation: State-of-the-Art and Future Research Opportunities’, Circulation, 125(23), pp. 2933–2943. Available at: 10.1161/CIRCULATIONAHA.111.069450.

Schmidt, C. et al. (2014) ‘Inhibition of cardiac two-pore-domain K+ (K2P) channels--an emerging antiarrhythmic concept’, European Journal of Pharmacology, 738, pp. 250–255. Available at: 10.1016/j.ejphar.2014.05.056.

Schmidt, C. et al. (2015) ‘Upregulation of K(2P)3.1 K+ Current Causes Action Potential Shortening in Patients With Chronic Atrial Fibrillation’, Circulation, 132(2), pp. 82–92. Available at: 10.1161/CIRCULATIONAHA.114.012657.

Silva Cunha, P., et al. (2024) ‘The impact of atrial voltage and conduction velocity phenotypes on atrial fibrillation recurrence’, Frontiers in Cardiovascular Medicine, 11, p. 1427841. Available at: 10.3389/fcvm.2024.1427841.

Skibsbye, L. et al. (2014) ‘Small-conductance calcium-activated potassium (SK) channels contribute to action potential repolarization in human atria’, Cardiovascular Research, 103(1), pp. 156–167. Available at: 10.1093/cvr/cvu121.

Smith, C.E.R., Ni, H. and Grandi, E. (2025) ‘Sex Differences in Electrophysiology and Calcium Handling in Atrial Health and Fibrillation’, Annual Review of Physiology, 87(Volume 87, 2025), pp. 1–24. Available at: 10.1146/annurev-physiol-022724-104938.

Sobie, E.A. (2009) ‘Parameter sensitivity analysis in electrophysiological models using multivariable regression’, Biophysical Journal, 96(4), pp. 1264–1274. Available at: 10.1016/j.bpj.2008.10.056.

Sossalla, S. et al. (2010) ‘Altered Na+Currents in Atrial Fibrillation’, Journal of the American College of Cardiology, 55(21), pp. 2330–2342. Available at: 10.1016/j.jacc.2009.12.055.

Streur, M. (2019) ‘Atrial Fibrillation Symptom Perception’, The journal for nurse practitioners : JNP, 15(1), pp. 60–64. Available at: 10.1016/j.nurpra.2018.08.015.

Thibault, S. et al. (2022) ‘Connexin Lateralization Contributes to Male Susceptibility to Atrial Fibrillation’, International Journal of Molecular Sciences, 23(18), p. 10696. Available at: 10.3390/ijms231810696.

Tobb, K., Kocher, M. and Bullock-Palmer, R.P. (2022) ‘Underrepresentation of women in cardiovascular trials-it is time to shatter this glass ceiling’, American Heart Journal Plus: Cardiology Research and Practice, 13, p. 100109. Available at: 10.1016/j.ahjo.2022.100109.

Tsai, W.-C. et al. (2016) ‘Hormone replacement therapy and risk of atrial fibrillation in Taiwanese menopause women: A nationwide cohort study’, Scientific Reports, 6, p. 24132. Available at: 10.1038/srep24132.

Tsuneda, T. et al. (2009) ‘Deficiency of testosterone associates with the substrate of atrial fibrillation in the rat model’, Journal of Cardiovascular Electrophysiology, 20(9), pp. 1055–1060. Available at: 10.1111/j.1540-8167.2009.01474.x.

Van Wagoner, D.R. et al. (1999) ‘Atrial L-Type Ca2+ Currents and Human Atrial Fibrillation’, Circulation Research, 85(5), pp. 428–436. Available at: 10.1161/01.RES.85.5.428.

Watanabe, R. et al. (2022) ‘Different Determinants of the Recurrence of Atrial Fibrillation and Adverse Clinical Events in the Mid-Term Period After Atrial Fibrillation Ablation’, Circulation Journal: Official Journal of the Japanese Circulation Society, 86(2), pp. 233–242. Available at: 10.1253/circj.CJ-21-0326.

Wettwer, E. et al. (2013) ‘The new antiarrhythmic drug vernakalant: ex vivo study of human atrial tissue from sinus rhythm and chronic atrial fibrillation’, Cardiovascular Research, 98(1), pp. 145–154. Available at: 10.1093/cvr/cvt006.

Wong, J.A. et al. (2017) ‘Menopausal age, postmenopausal hormone therapy and incident atrial fibrillation’, Heart (British Cardiac Society), 103(24), pp. 1954–1961. Available at: 10.1136/heartjnl-2016-311002.

Yang, P. et al. (2016) ‘In silico prediction of drug therapy in catecholaminergic polymorphic ventricular tachycardia’, The Journal of Physiology, 594(3), pp. 567–593. Available at: 10.1113/JP271282.

Yue, X. et al. (2017) ‘Heterogeneity of transverse-axial tubule system in mouse atria: Remodeling in atrial-specific Na+-Ca2+ exchanger knockout mice’, Journal of Molecular and Cellular Cardiology, 108, pp. 50–60. Available at: 10.1016/j.yjmcc.2017.05.008.

Zhang, X. et al. (2024) ‘Enhanced Ca2+-Driven Arrhythmogenic Events in Female Patients With Atrial Fibrillation’, JACC: Clinical Electrophysiology, 10(11), pp. 2371–2391. Available at: 10.1016/j.jacep.2024.07.020.

